# Novel antibody language model accelerates IgG screening and design for broad-spectrum antiviral therapy

**DOI:** 10.1101/2024.03.01.582176

**Authors:** Hannah Faisal Almubarak, Wuwei Tan, Andrew D. Hoffmann, Yuanfei Sun, Juncheng Wei, Lamiaa El-Shennawy, Joshua R. Squires, Nurmaa K. Dashzeveg, Brooke Simonton, Yuzhi Jia, Radhika Iyer, Yanan Xu, Vlad Nicolaescu, Derek Elli, Glenn C. Randall, Matthew J. Schipma, Suchitra Swaminathan, Michael G. Ison, Huiping Liu, Deyu Fang, Yang Shen

**Author notes:** Correspondence to: Yang Shen, PhD, Texas A&M University, College Station, TX 77843. Deyu Fang, MD, PhD, Northwestern University, 300 E Chicago St, Chicago, IL 60611. Huiping Liu, MD, PhD, Northwestern University, 303 E Superior St, Chicago, IL 60611. Co-first authors with equal contributions (Hannah Faisal Almubarak, Wuwei Tan, Andrew D. Hoffmann, Yuanfei Sun, Juncheng Wei).

## Abstract

Therapeutic antibodies have become one of the most influential therapeutics in modern medicine to fight against infectious pathogens, cancer, and many other diseases. However, experimental screening for highly efficacious targeting antibodies is labor-intensive and of high cost, which is exacerbated by evolving antigen targets under selective pressure such as fast-mutating viral variants. As a proof-of-concept, we developed a machine learning-assisted antibody generation pipeline AbGen that greatly accelerates the screening and re-design of immunoglobulins G (IgGs) against a broad spectrum of SARS-CoV-2 coronavirus variant strains. Our AbGen centers around a novel antibody language model (AbLM) that is pretrained on 12 million generic protein domain sequences and fine-tuned on 4,000+ paired VH-VL sequences, with IgG-specific CDR-masking and VH-VL cross-attention. AbLM provides a latent space of IgG sequence embeddings for AbGen, including (a) landscapes of IgGs’ activities in neutralizing the wild-type virus are analyzed through structure prediction for IgG and IgG-antigen (viral protein spike’s receptor binding domain, RBD) interactions; and (b) landscapes of IgGs’ susceptibility in neutralizing variant viruses are predicted through Gaussian process regression, despite that as few as 14 clinical antibodies’ responses to variants of concern are available. The AbGen pipeline was applied to over 1300 IgG sequences we collected from RBD-binding B cells of convalescent patients. With experimental validations, AbGen efficiently prioritized IgG candidates against a broad spectrum of viral variants (wildtype, Delta, and Omicron), preventing the infection of host cells *in vitro* and hACE2 transgenic mice *in vivo*. Compared to other existing protein language models that require 10-100 times more model parameters, AbLM improved the precision from around 50% to 75% to predict IgGs with low variant susceptibility. Furthermore, AbGen enables structure-based computational protein redesign for selected IgG clones with single amino acid substitutions at the RBD-binding interface that doubled the IgG blockade efficacy for one of the severe, therapy-resistant strains - Delta (B.1.617). Our work expedites applications of artificial intelligence in antibody screen and re- design combining data-driven protein language models and Kriging for antibody sequence analysis and activity prediction, in synergy with physics-driven protein docking and design for antibody-antigen interface analyses and functional optimization.

## Introduction

One of the most demanding challenges in medical care and clinical investigation is to fight against constantly evolving pathogens and abnormal cells (such as cancer) under selective therapeutic pressure. Antibodies have accounted for most of the top therapeutic sales impacting medical treatments for immune diseases and cancer over the past years (1–3). However, the traditional development of any therapeutic drugs including antibodies is time- and labor-intensive, requiring a large scale of experimental screening and validation. Many antibody screening strategies for neutralization efficacy such as phage display, ribosome display, and mammalian cell surface display are relatively of low efficiency (4). In this study, we set out to accelerate antibody development by integrating a novel antibody language model (AbLM) and antibody structure, interaction, and viral-neutralizing activity prediction with experimental validations into a pipeline (AbGen) for broad-spectrum antiviral therapy, utilizing the fast-mutating severe acute respiratory syndrome coronavirus 2 (SARS-CoV-2) as a targeting model.

SARS-CoV-2 enters host cells via its viral spike protein binding with the host cell receptor angiotensin-converting enzyme 2 (ACE2) (5, 6), which is highly expressed on the cell membrane in various human organs including the lungs, heart, and kidney (7). During the pandemic of coronavirus disease 2019 (COVID-19), neutralizing antibodies became one of the earliest approaches that rapidly developed and effectively treated the early phase of infections before a vaccine was developed and before the coronavirus evolved to evade neutralization (8–12). Most of the coronavirus neutralizing antibodies disrupt the spike interactions with human ACE2 thereby preventing viral entry for prophylactic and therapeutic applications (8–15).

The constantly-evolving viral sequences especially the variants of concern (Delta and Omicron) started to escape from neutralizing antibodies and vaccination immunity (16, 17), partially contributing to the death of nearly seven million people worldwide (18). For instance, the spike L452R and T478K mutations in the Delta variant, also known as B.1.617 lineage, are located at the periphery or the epitope region of the receptor binding domain (RBD) and are found to reduce antibody neutralizing activity (19, 20). The Omicron variant, which is characterized by the presence of around 32 mutations in the RBD, escapes most therapeutic neutralizing antibodies and largely vaccine-elicited antibodies (17).

To better prepare for any inevitable pandemics or epidemic diseases, rapid development and design of broad-spectrum antibodies would facilitate the early response against infectious pathogens (21). The emerging artificial intelligence has shown a great potential to expedite and transform the antibody production pipeline (21, 22). However, it is unclear whether and how the therapeutic development could be accelerated in low-data regimes, where data are very limited for epitope identification, structure identification, and variant responses, which curbs the power of data-hungry computational methods such as machine learning. Despite the critical barriers, we integrated experimental data, physics simulations, and machine learning into an antibody-screening and optimization platform. With limited experimental data, we built and validated such methods in predicting and ranking patient antibodies against broad coronavirus variants.

To improve the efficiency of experimental screening, we built our computational pipeline AbGen on a novel antibody language model, AbLM, through pretraining over twelve million generic protein domain sequences and finetuning over four thousand paired VH-VL sequences, with antibody-specific CDR masking and VH-VL cross-attention. No activity data was required for AbLM. In the latent space of IgG sequence embeddings suggested by AbLM, we first used physics-driven IgG antibody structure prediction and IgG-RBD docking to predict the IgG neutralization landscape against the wild-type (WT) coronaviral strain. Again no activity data was required here for predicting WT neutralization. Second, using known activity data of as few as 14 antibodies, we constructed Gaussian process regressors (23) in the latent space to predict the IgG neutralization landscape against the viral variants of concern. Furthermore, we used computational protein design with the input of protein-docking structural models to redesign IgGs for improved neutralization efficacy against the Delta variant. The screened or redesigned IgGs were validated in both *in vitro* and *in vivo* virus infection studies.

## Results

### Overview of the machine learning-accelerated antibody production platform

Using SARS-CoV-2 as a targeting model pathogen and the spike RBD as a bait antigen, we developed a low data regime-derived pipeline facilitating the screening and redesign of neutralizing antibodies (**Fig. 1a)**. To identify and produce anti-spike antibodies, we first collect blood specimens from 42 patients recovered from early COVID-19 as described (24), flow sorted the RBD-bound IgM memory B cells for subsequent single-cell VDJ sequencing via the 5’ 10X genomics platform, and retrieved 1376 heavy-light chain pairs of IgG antibody sequences.

**Figure 1:**
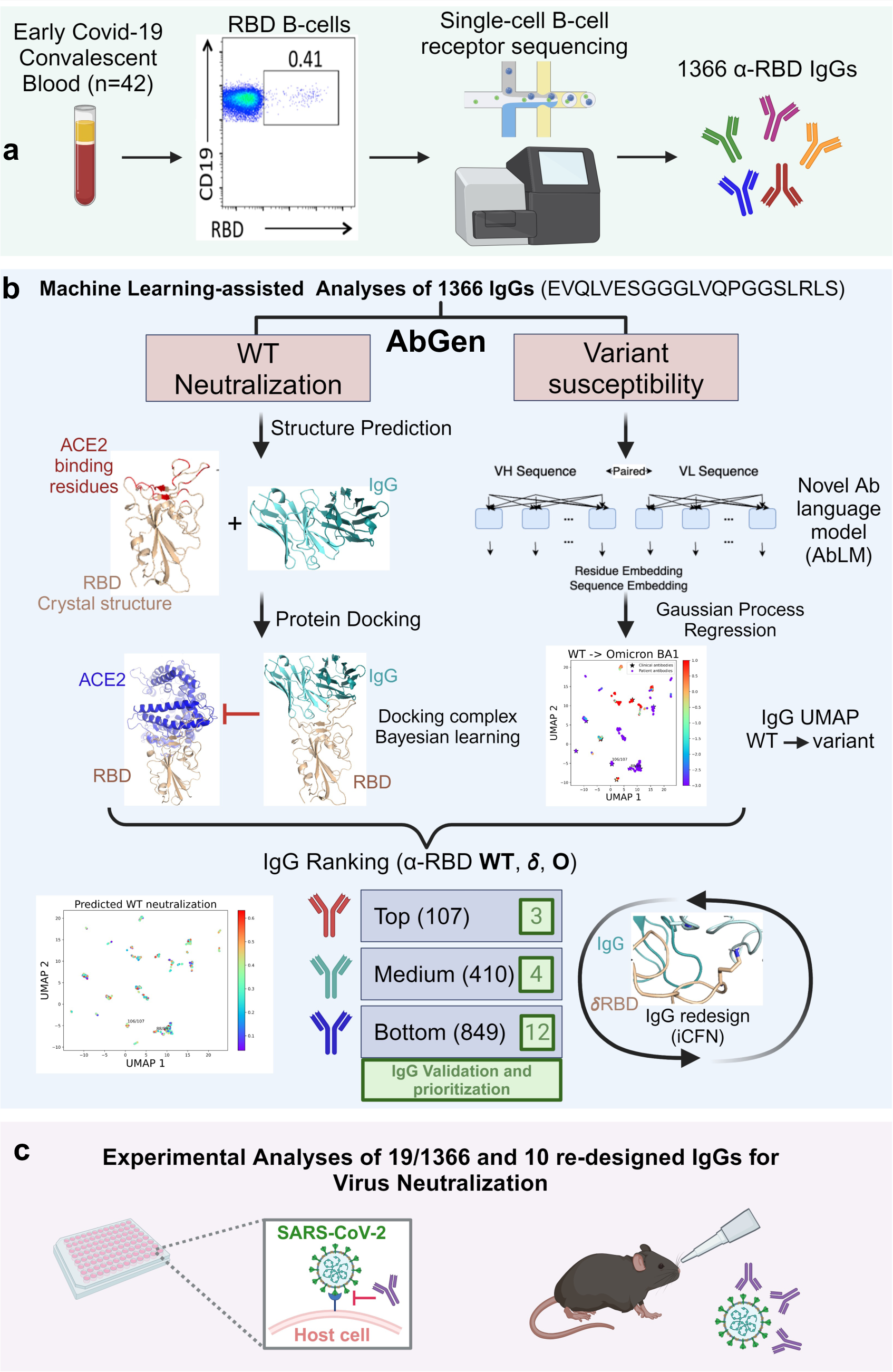
Schematic illustration of our study flow and the AbGen pipeline. **a.** Peripheral blood samples were collected from 42 convalescent Covid-19 patients and RBD+ memory B-cells were sorted and underwent single-cell VDJ sequencing using the 10x Genomics platform. **b.** A total of 1366 a-RBD antibody sequences were retrieved and included in our machine-learning model to predict antibodies with high SARS-CoV2 neutralization capacities. **c.** Prioritized antibodies were then tested for their ability to neutralize broad-spectrum SARS-CoV-2 in both in-vitro and in-vivo settings.

Facing the low data regimes where activity data are available for none or few antibodies, we harnessed two distinct approaches, namely physics-driven protein docking and data-driven machine learning, for the prediction of antibodies’ effectiveness in virus neutralization (**Fig. 1b** with details in **Fig. S1**). We first predicted the antibody structure of each sequence derived from convalescent patient samples. Subsequently, leveraging both the predicted antibody structures and the crystal structure of the spike RBD, we generated multiple docking configurations of antibody-RBD structures. Later, we refined a confidence score for each IgG docking structure, which is predicted as a coverage fraction or blockade portion of the RBD residues for ACE2 binding, serving as a physics-driven predictor for WT virus neutralization. Higher coverage of the ACE2 binding residues would predict better neutralization. This structure-based protein docking approach has been previously validated for its accuracy in modeling antigen-antibody complex structures (25).

To predict neutralization against variants of concern such as Delta and Omicron strains, we calculated the covariances (variograms) to represent dissimilarities or distances among all sequenced antibodies and clinically used antibodies, using the latent space learned in an antibody language model. Then, we predicted the robustness of sequenced antibodies to neutralize variants using Kriging (Gaussian process regression) (23), with the input of the variograms above and the IC_50_ fold changes of 14 clinical antibodies from WT to variants. This sequence-based antibody language model is novel in tailoring pretrained protein language models for paired VH-VL chains and later shown to outperform other language models in variant response prediction.

Based on the confidence score against the WT virus and the robustness predictions against viral variants, the IgG ranking facilitated IgG prioritization and re-design for further experimental validation. After machine learning-assisted neutralization prediction classified all sequenced IgG antibodies into distinct categories of top, medium, and low confidence (or priorities), we cloned 19 randomly picked IgG pairs of heavy and light chains spanning these categories and a few computationally redesigned antibodies for experimental analyses and functional neutralization against SARS-CoV-2 infections *in vitro* and *in vivo* (**Fig. 1c**).

### Diversity of sequenced antiviral antibodies from human B cells

After obtaining 1376 heavy-light chain pairs of IgG antibodies using the CellRanger platform (10X genomics), we identified the individual V, D and J genes (sequences) for each antibody as well as the type of light chain (Kappa or Lambda). Frequency analyses of individual V and J genes revealed a non-even distribution, as certain V and J genes were more common with 60 to 600 counts than others with few counts (**Fig. S2a-b**). D genes were not detected in the light chains and were rarely seen in the heavy chain (**Fig. S2c**). The relative diversity of V genes was identified with the J genes skewed toward a few sequences, such as IGHJ4 of over 600 counts, suggesting that our sample group of antibody candidates share general properties of IgGs with distinct specificity of how each IgG recognizes and binds to spike-RBD. We detected a few C genes in the heavy chains and Lambda light chains but not within the Kappa chains (**Fig. S2d**). Every CDR3 sequence within the H chains was identified once, similar to most of the CDR3 sequences in K and L chains with only a small portion repeating 2-10 times (**Fig. S2e, f**).

Since it is labor-intensive and inefficient to experimentally test 1000+ IgG candidates for functional efficacy tests, we sought to utilize physics-driven protein docking and data-driven machine learning for computational predictions that accelerate the experimental screen and redesign of efficacious neutralization antibodies.

### Antibody language model provides a latent space for IgG analysis and design

To analyze the sequence distribution of IgG candidates at a large scale without time-consuming sequence alignment, and to further improve prediction of antibody activities, we designed and trained a novel antibody language model (AbLM) (**Fig. 2a)**. AbLM utilizes readily available, unlabeled protein sequence data for pretraining, paired antibody heavy and light chain sequences for finetuning, and CDR-informed masking (see details in Methods). For the convalescent IgG antibodies our sequenced as well as 14 clinically used COVID antibodies, we embedded their variable region sequences into 1536-dimensional vectors (768 dimension for either heavy or light chain) using the paired sequence encoders of AbLM and visualized their 2D distributions using Uniform Manifold Approximation and Projection (UMAP) (26) (**Fig. 2b**). We noticed that the 14 clinically proven antibodies were clustered with (or “covered” by) some convalescent IgG clusters, which suggests the clinical potential to prioritize and improve selected IgG antibodies. As demonstrated in the subsequent subsections, AbLM provided a capability to harness the potential.

**Figure 2.**
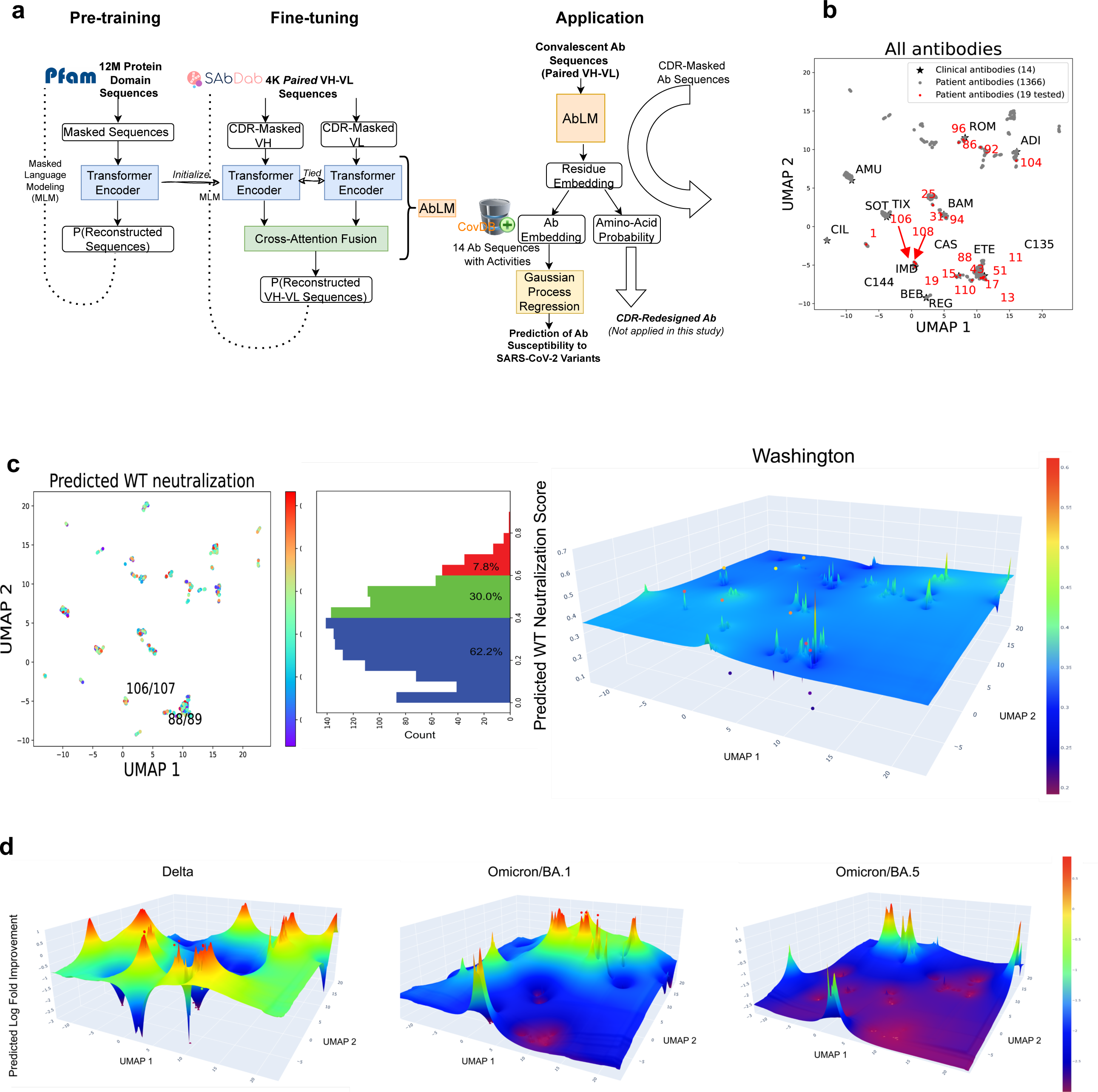
Our novel Antibody Language Model (AbLM) embeds IgG sequences in a latent space for sequence analysis and susceptibility prediction. **(a)** Architecture, training, and application of our novel antibody language model, AbLM. **(b)** Clusters of 1366 patient-derived convalescent antibodies along with 14 clinical antibodies, based on the sequence embeddings from AbLM and dimensionality reduction using UMAP. 19 antibodies highlighted in red were functionally tested. **(c)** Predicted 2D and 3D landscape of patient antibodies’ neutralization of wild type (a confidence score between 0 and 1), along with a 1D histogram in between, using antibody-antigen structure predictions. **(d)** Predicted 3D landscape of patient antibodies’ robustness to variants (log fold improvement in neutralization compared to that of the wild type / Washington, ranging from -3 to +1), using Gaussian Process Regression (Kriging) based on 14 clinical antibodies’ known variant responses.

On the UMAP, the randomly selected 19 IgG candidates (in red color) for experimental validation showed diversity among themselves and coverage of the cluster sets of over 1000 antibodies (**Fig. 2b**). Based on the AbLM analysis, IgG 106/107, named after H/L indices, resembled the sequence of clinical antibodies C144 (8) and IMD (27) (Imdevimab or REGN10987) whose neutralization mechanism involves direct competition with ACE2; whereas IgG 88/89 resembled CAS (27) (Casivirimab or REGN10933). Sequence alignment proved such resemblance that 106/107 and C144 share 11 of 12 CDR3 amino acids in the light chain (CSSYTSSST*G*VF versus CSSYTSSST*R*VF), although C144 has a much longer CDR3 sequence than 106/107 (14 amino acids for 106/107 versus 23 amino acids for C144) (**Supp. Fig. S3a**).

### Physics-driven prediction of IgG structures, RBD interactions and WT viral neutralization landscape

To assess mechanistically and virtually the neutralization capabilities of these uncharacterized IgG candidates, we established a physics-driven prediction pipeline (**Fig. S1b**). It simulates IgG apo-structures (H/L pairs) via AbodyBuilder-ML (28) and predicts IgG halo-structures bound to spike RBD, using HADDOCK (25) for initial docking and employing Bayesian Active Learning (BAL) (29) for refinement and uncertainty quantification. We hypothesized that an IgG antibody of higher neutralization capability blocks more competitively the viral RBD’s binding to the entry receptor ACE2 as one neutralization mechanism. Therefore, we used the IgG-competing portion of the ACE2-binding residues of RBD (0.0-1.0) as a confidence indicator to predict the IgG’s neutralization against the WT virus (**Supp. Table 1** [Tab 1 “All IgGs (WT and variant pred.)”]). The predicted IgG groups are visualized as scatter points and the smoothened neutralization landscape in the latent space (**Fig. 2c**). The majority, 849 (>62%) of the 1366 IgG candidates (1376 excluding 10 with failed apo-structure prediction) were predicted with a confidence score below 0.4, implying a low neutralization potential against the WT thus a low priority. 107 IgGs (7.8%) were predicted with the scores above 0.6 (**Fig. 2c**) and these high-priority antibodies were dispersed in the latent space (**Fig. S3a**), including regions not covered by the 14 clinical antibodies and suggesting novel sequences of clinical application potentials.

### Data-driven prediction of neutralization profiles against the viral variants in low-data regimes

To predict the neutralization profiles of the uncharacterized IgG candidates, we followed an early response scenario of “few-shot learning” by selecting a few IgGs for experimental characterization and providing a few labeled data for machine learning. The IgG selection can be based on the diversity (clusters covering the latent space) and the predicted WT neutralization. Before any of our IgG candidates was tested, we emulated the selection process using the 14 clinical antibodies that satisfied the criteria and had been experimentally characterized for many SARS-CoV-2 variants. As machine learning especially large deep learning models often demand big data that is unavailable in the early response scenario or other disease settings, we chose Gaussian process regressor (Kriging), the best linear unbiased estimator, in combination with a latent space from a protein language model, to predict variant-response robustness (the IC_50_ log-fold improvement of each IgG candidate in neutralizing variants versus WT) based on just the 14 clinical antibody profiles.

For all 1366 IgG candidates, we visualized the predicted robustness landscape against Delta and Omicron BA1/BA5 variants (evolving variants compared to the WT infection the convalescent patients experienced) in the latent space (**Fig. 2d** and **Supp. Fig. S3b** with data provided in the **Supp. Table 1** [Tab 1 “All IgGs (WT and variant pred.)”]). We observed a few regions enriched in better robustness for Delta variant (warmer colors), such as those around IgG 106/107 predicted to resemble nearby IMD (Imdevimab/REGN10987) and neutralize the Delta variant well (but losing robustness against the Omicron BA1 or BA5 variants). The prediction revealed an overall cooler colored landscape against the Omicron variants versus the Delta variants, echoing the observations from other studies (17, 30) that the Omicron variants are more prone than the Delta variant to elicit antibody escape.

### Functional tests of IgG candidates in neutralizing RBD binding and viral infections

To assess our predictions of IgG candidates in neutralizing the WT and variant strains of SARS-CoV-2, we *randomly* selected 19 IgG antibodies for experimental tests, including three [88/89, 106/107, and 94/95] in the high priority class, four in the medium priority class, and 12 in the low priority class against WT (**Fig. 3a** and **Supp. Table 1** [Tab 2 “Validated IgGs (WT pred.)”]). Following heavy and light chain cloning, the plasmid DNA was transfected into HEK293T cells for IgG overexpression, and the IgGs were purified for functional analyses in blocking RBD binding to host cells and neutralizing live viral infections.

**Figure 3.**
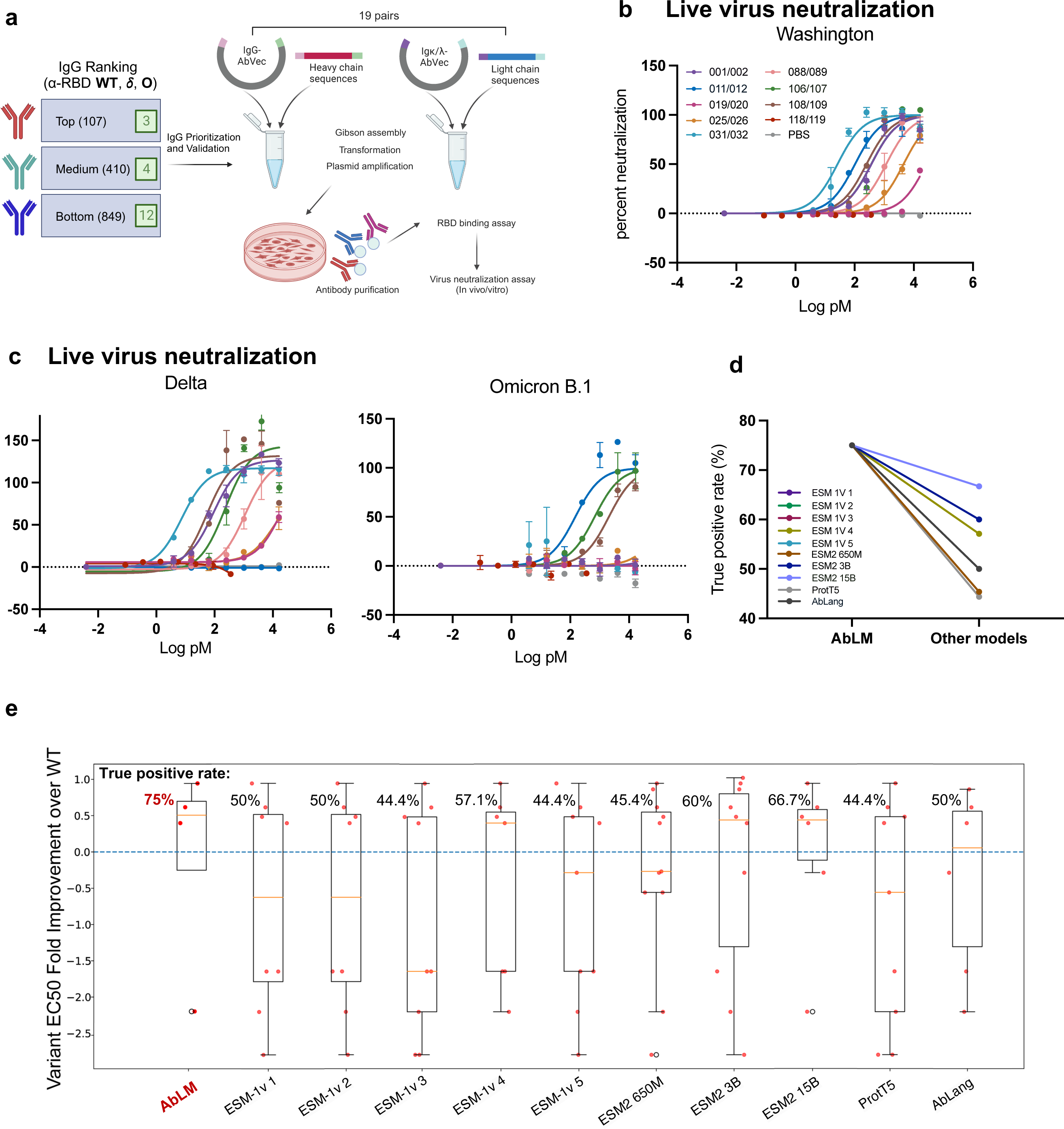
Experimental tests of antibody neutralization against SARS-CoV-2 WT and variants. (a) Schematic depicting experimental workflow. Variable regions of the heavy and light chains of the prioritized antibodies were cloned into AbVec IgG vectors followed by transfection into 293T cells. Purified antibodies were then tested for their ability to neutralize RBD in an ACE-2 cell-binding assay as well as live SARS- CoV2 neutralization. (b, c) Live SARS-CoV-2 virus neutralization by prioritized anti-RBD antibodies. Viability of A549 cells overexpressing human ACE2 was evaluated 72-96 hours post viral infection. GraphPad Prism 9.3.1 was used to calculate EC50s (d) Box plots of antibodies tested antibodies’ wild-type neutralization (binding inhibition IC50 and viral neutralization EC50) split into three predicted to be of to (e) Box plots of tested antibodies’ variant response (log fold improvement in viral neutralization EC50 compared to the wild type) split into four antibody-variant cases predicted to be robust and eighteen predicted to be susceptible.

With flow cytometry-based measurement of RBD-binding to human ACE2 expressing cells (31), we evaluated the IgG abilities in blocking the RBD-binding to cellular ACE2 receptor (binding inhibition IC_50_) (**Supp. Fig. S4a**). Seven out of 19 randomly selected then tested IgGs (001/002, 019/020, 025/026, 031/032, 088/089, 106/107 and 108/109) inhibited the RBD binding to ACE2^+^ HEK293 cells in a dose dependent manner (**Supp. Fig. S4a** and **Supp. Table 2**). Among the seven neutralizing IgGs, two were extremely potent (106/107 and 31/32) with IC50 < 0.5 nM, both of which confidence scores are among the top (**Fig. 2**). The rest 12 antibodies did not show any effective neutralization activity at the concentration tested (**Supp. Table 2**), all of which are largely consistent with their low confidence scores (**Fig. 2c**).

Consistently, nearly half of the IgG antibodies had neutralizing effects at serially diluted concentrations (up to 16 nM) on live SARS-CoV-2 infection to ACE2 overexpressing A549 cells (**Fig. 3b**). Specifically, the IgG clones 106/107, 011/012 and 031/032 showed the lowest EC_50_ values against the WT Washington strain, demonstrating the strongest neutralizing effects (**Fig. 3b**). Interestingly, although the IgGs were derived from the COVID-19 patients infected with the WT Washington strain (before the variants of concern emerged and widespread), many of the IgGs possessed neutralizing capacities against the Delta and Omicron variants as well. In fact, three of the clones (001/002, 031/032, and 108/109) were about ten times more effective in neutralizing the Delta variant as compared to the wildtype (**Fig. 3c**, **Fig. S4b** and **Supp. Table 3**).

These experimental data validated our computational pipelines’ predictive power in screening and prioritizing potent and broad-spectrum antibodies, as detailed as follows.

First, independent of IgG activity data, our physics-driven confidence prediction for IgGs in WT neutralization, based on the predicted blocking portion of ACE2-binding RBD residues, showed robust prioritization of WT neutralizing antibodies (**Fig. S5a** and **Fig. 2c**). The first two of the top three IgGs with high-confidence score priority (88/89 and 106/107) showed high potencies and efficacies in neutralizing RBD-binding and viral infection, representing a 67% success rate compared to 37% (7/19) from a random subset of antibodies. Conversely, among the twelve predicted low-priority antibodies, 75% and 67% were proven to have no activity in neutralizing RBD-binding and viral infection, respectively (**Fig. S5a**).

Second, using data from as few as 14 antibodies (**Supp. Fig. S6**), our data-driven prediction profiles of antibodies on variant neutralization also showed agreement with the experimental inhibition of variant infection in differentiating narrow- and broad-spectrum antibodies (**Fig. 3d-e** and **Fig. 2d**). Among 22 experimentally validated antibodies against variant infections, 3 out of 4 predicted to have improved neutralization efficacy (better robustness compared to the WT virus) were validated with a success rate of 75% (3/4), whereas 14 of the rest 18 predicted to lose efficacy were also validated with a success rate of 78% (14/18) (**Fig. 3e**, and **Supp. Fig. S5b**).

The success rates from our antibody language model, especially that for robust responses (75%), significantly outperformed those from state-of-the-art pretrained language models, including all five versions of ESM-1 (32) (44%–57% for robustness), three largest versions of ESM-2 (33) (45%–67%), ProtT5 (34) (44%), and an antibody language model AbLang (35) (50%), as shown in **Fig. 3d,e** and **Supp. Fig. S5b**. Compared to the next best performer ESM-2 with 15 billion parameters, our AbLM with 92 million parameters (163-times smaller) is accurate yet lightweight, thanks to antibody-inspired novelty including pairing VH-VL encoders with cross attention and training them with CDR-masking.

### Redesign of a prioritized antibody improves neutralizing the Delta variant

Given the demonstrated ability of our platform to prioritize potent and broad-spectrum antibodies, we next tested its ability to (re)design prioritized IgGs for better variant neutralization profiles. We picked IgG 106/107 that was a top candidate of our predictions and verified with its neutralizing efficacy against WT and the Delta variant. Based on predicted IgG-RBD structures, we used a physics principle-driven multistate protein design program, iCFN (36) to computationally redesign IgG 106/107 to improve its binding to the Delta variant RBD without losing much affinity to the WT RBD. Ten single amino-acid substitutions near suggested binding sites of T478K were proposed, including three at the heavy chain (all on F66) and seven at the light chain (four on Y38, two on Y55 and one on D56) (**Fig. 4a** and **Supp. Fig. S7**). All four designs at Y38 stood out with a higher confidence score due to strongly improved electrostatic interactions, half being large-to-small apolar substitutions (Y38A and Y38G) and the other half hydrophobic-to-polar (Y38S and Y38Q).

**Figure 4.**
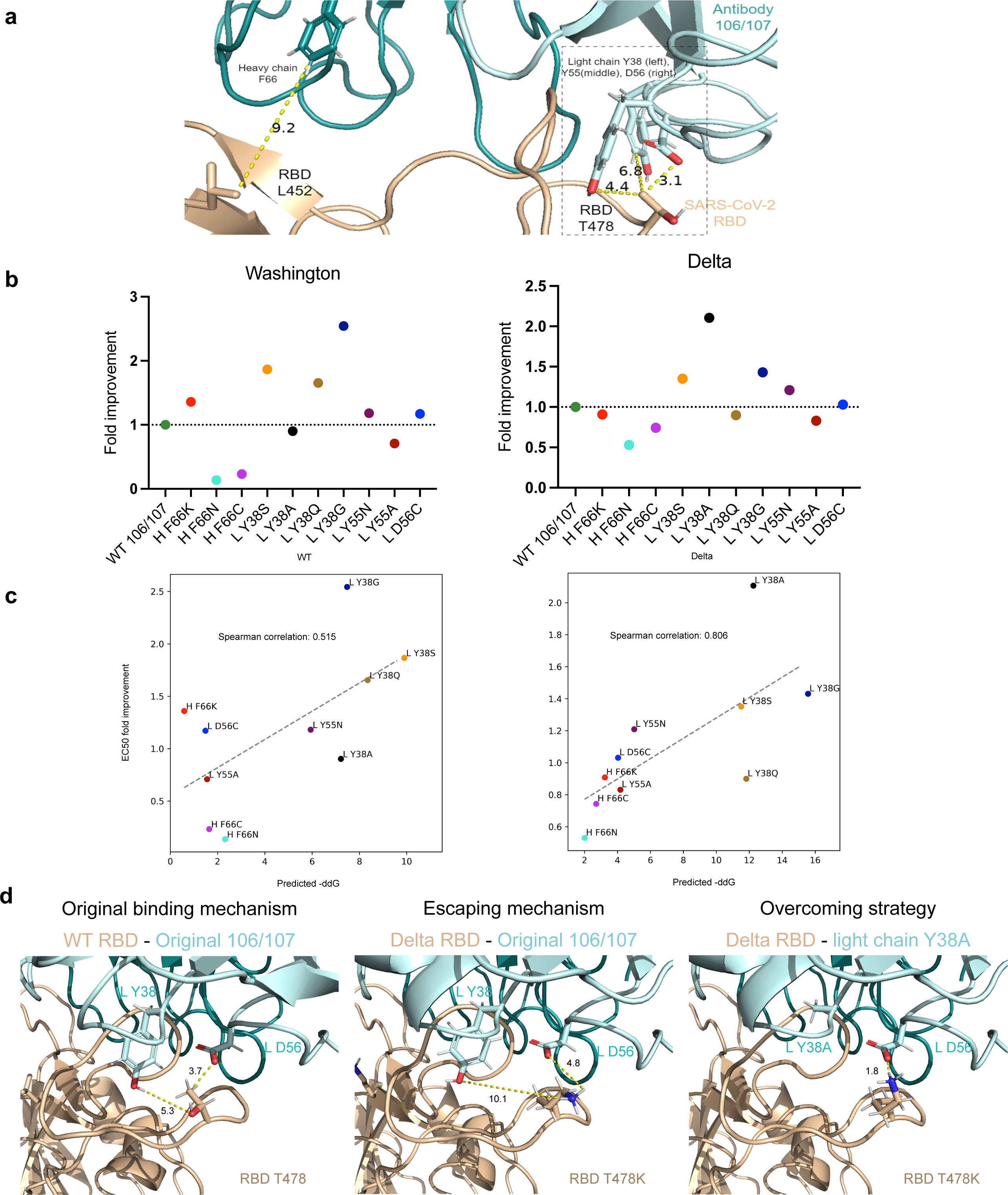
Antibody 106/107 redesign with improved efficacy against delta variant. (a) Visualization of the binding interface between the spike RBD (in wheat cartoon), including two positions mutated in the Delta variant (L452 and T478), and the patient antibody 106/107 (in blue cartoons). Four positions on the antibody were selected for redesign and shown in sticks, including F66 of the heavy chain (in darker blue) and Y38, Y55, and D56 of the light chain (in lighter blue). (b) Live SARS-CoV-2 virus neutralization by our redesigned antibodies. Data showing fold-improvement of EC50 for the redesigned antibodies compared to the WT 106/107 antibody. Viability of A549 cells overexpressing human ACE2 was evaluated 72-96 hours post viral infection. GraphPad Prism 9.3.1 was used to calculate EC50s (c) Computational designed single amino-acid substitutions of antibody 106/107: computationally-predicted binding-energy improvements (-ddG) showed high correlations with experimentally-measured EC50 fold improvements, whether tested against WT (Washington) or the Delta variant. (d) Computationally predicted 3D structures of RBD-Ig complexes suggested the WT-neutralizing mechanism of the antibody 106/107 (where Y38(L) and D56(L) interacted with T478 on the RBD wild type), the antibody-escaping mechanism of the Delta variant (where Y38(L) and D56(L) lost interactions with K478 on the RBD Delta variant and Y38(L) paid more desolvation penalty), and the variant-neutralization enhancing mechanism (where A38(L) paid almost no desolvation penalty).

We cloned ten re-designed IgG 106/107m antibodies and measured their neutralization abilities against live viruses of the wild type and Delta variant. Strikingly, three of the four top-ranked designs, all at Y38 residue including Y38A, Y38S and Y38G (not Y38Q) showed most improved neutralization against the Delta variant, with Y38A of over 2-fold increase in neutralization efficacy (**Fig. 4b and Supp. Fig. S8**). Importantly, their neutralization activities for the WT virus were not significantly impaired, which is consistent with the intended computational design. Our principle-driven protein design led to energy scores with positive to strongly positive ranking performances: the Spearman’s rank correlation coefficient between the predicted binding energy changes and the measured pEC50 values for the 10 designs was 0.515 and 0.806 for wild type and the Delta variant, respectively (**Fig. 4c**). It also led to mechanistic explanations of Y38A’s gained binding to the Delta RBD: T478K in the Delta variant may cause Y38 of the antibody 106/107 to be buried from the solvent and pay desolvation penalty to the binding affinity (**Fig. 4d** middle panel), whereas the substitution to a small hydrophobic residue Y38A (as well as Y38G) in the antibody 106/107 can reduce such penalty (**Fig. 4d** right panel). With the proof of concept from single substitution designs, we anticipate that higher-order redesigns or even *de novo* designs would further improve the broad-spectrum neutralization profile.

### Prioritized IgG candidates show anti-infection in transgenic mice

We next evaluated the therapeutic efficacy of two best neutralizing IgG candidates (106/107 and 011/012) prioritized by our integrated computational and experimental platform in their ability to prevent multi-strain SARS-CoV-2 infection in the well-established hACE2 transgenic mouse model *in vivo* (**Fig. 5a**). Antibodies were administered to mice by nasal inhalation to determine their effects on preventing viral infection of the three viral strains (WT Washington, Delta, and Omicron). In concordance with the *in-vitro* cell survival after viral infections, both antibodies prevented weight loss and health deterioration in mice infected with the Washington strain, whereas only IgG 106/107, but not IgG 011/012, prevented the mice from Delta variant infection-caused weight loss and clinical symptoms (**Fig. 5b, c**). Furthermore, the viral titers in the lungs of infected mice were significantly lower after the 106/107 antibody treatment which also neutralized the viral variants, consistent with the live virus neutralization results *in vitro* (**Fig. 5d**).

**Figure 5.**
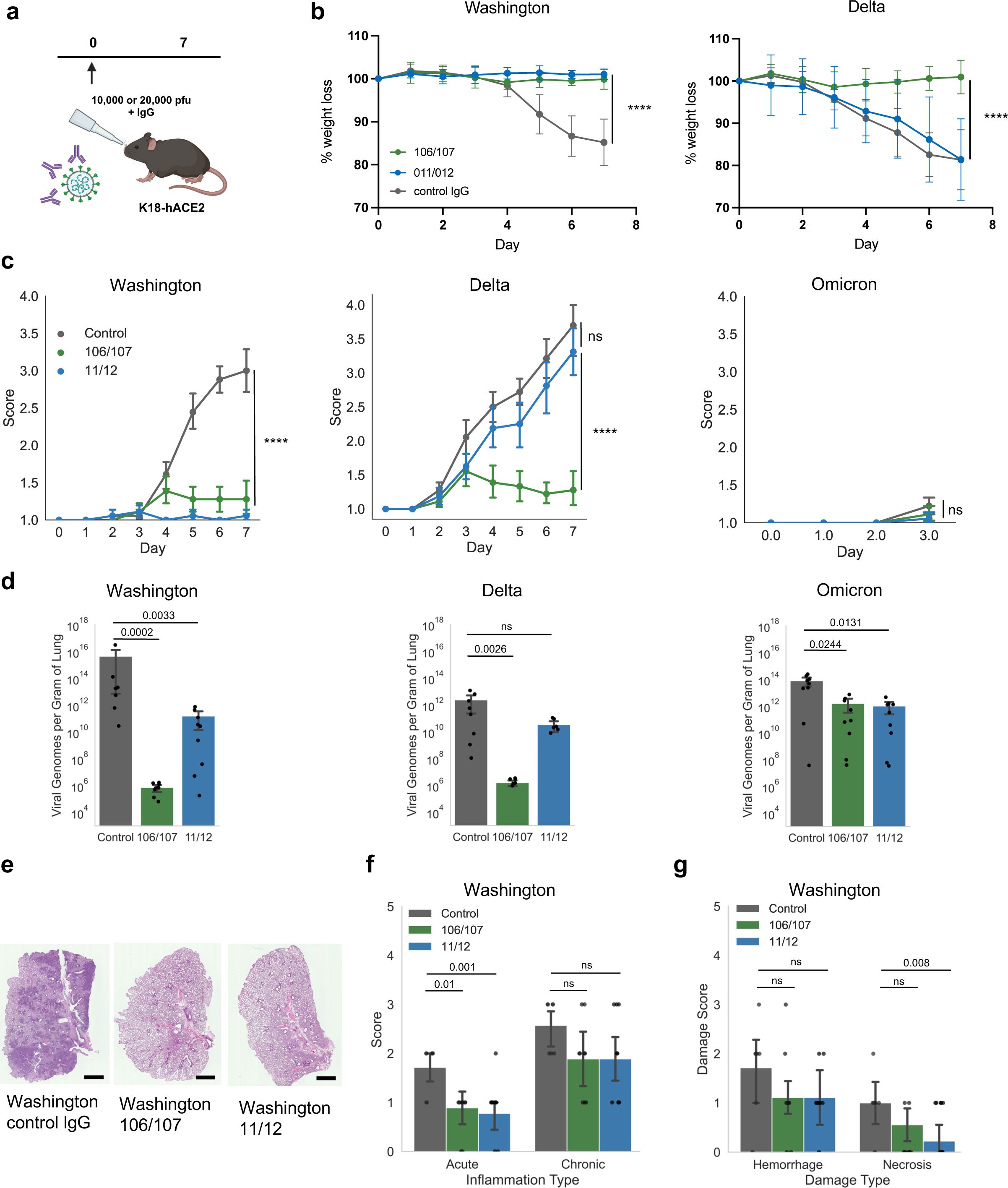
Antibody neutralization and inhibition of broad strain infections in hACE2 transgenic mice. (a) Schematic of viral neutralization and infection design. SARS-CoV-2 viral strains and select antibodies (1.2 ug IgG for Washington and Delta or 8.5ug IgG for Omicron) were administered intranasally into K18-hACE2 mice. Animals were monitored for health twice daily and weighed once per day. Seven days post infection (3 days for Omicron), mice were euthanized, and lungs were analyzed for viral load and pathohistological characteristics. (b) Percent weight loss of K18-hACE2 mice post viral infection (n= 9). (c) Clinical score post SARS-CoV-2 infection. Larger clinical scores indicate increased disease severity and reduced physical fitness (n= 9). (d) Lung genomic viral load on day 7 (Washington and Delta) or day 3 (Omicron) post antibody neutralization and viral infection measured by qRT-PCR (n= 9). (e) Representative H&E images of mice lungs at experimental endpoint. Error bar = 1mm. (f) Lung Acute and chronic inflammation scores based on histopathological analysis of H&E-stained slides (n= 9). (g) Alveolar hemorrhage and necrosis scores based on histopathological analysis of H&E-stained slides (n= 9). Statistical significance was tested with unpaired t-tests (b, c, f and g) or Mann Whitney test (d). ns means non-significant difference.

Notably, while the low pathogenic Omicron variant did not cause severe disease in mice, treatment with either IgG 106/107 or IgG 011/012 resulted in lower levels of viral titers in the lungs of the infected animals (versus the isotype IgG control group) (**Fig. 5c, d**). As expected, Omicron infection did not cause significant changes in clinical scores and lung histology even in the control mice without candidate treatment (**Fig. 5c, d**, and **Supp. Fig. S9, S10**). Histology analyses of the lungs of infected mice reveal that the 106/107 and 011/012 neutralizing IgGs were able to reduce acute lung inflammation following infection with the Washington and Delta strains, albeit no significant effect on chronic inflammation (**Fig. 5e, 5f**, and **Supp. Fig. S9, S10**). Additionally, the IgG 011/012 antibody treatment significantly inhibited the necrosis in the lungs of mice infected with the Washington as compared to the control IgG (**Fig. 5g**). These data demonstrated the IgG prioritization strategies in pre-clinical studies.

## Discussions and Conclusions

Therapeutic antibody development faces the dual challenges of expensive experimental screening and constantly evolving targets under selective pressure. Using SARS-CoV-2 as a testing model, our extensive RBD-specific IgG sequencing on a sizable scale with COVID-19 patients in the early stages of the pandemic, reveals that a significant number of effective clones against the WT virus, lose effectiveness against the variants of concern, such as Delta and Omicron. Significantly, our computational analysis pinpoints certain IgG antibody clones with potential enhancements in blocking variant infections in ACE2^+^ cells, as confirmed through both *in vitro* and *in vivo* virus infection studies. These findings offer a potential explanation for why certain COVID-19 patients exhibit greater resistance to reinfection by SARS-CoV-2 variants compared to others, as frequently observed (37).

Furthermore, we have demonstrated that replacement of a single residue spin in the IgG core for RBD binding with a smaller and hydrophobic alanine (Y38A) dramatically improves its neutralization activity against the Delta variant. This unexpected discovery also provides a rationale for us to harness the computational power in accelerated virtual IgG screening and re- design for their use to treat/prevent emerging new pathogen variants or mutating oncogenic targets, which are urgent clinical demand. Over the last few years, while the convalescent plasma treatment has been proven to be effective in treating COVID-19 patients [30], almost all licensed monoclonal antibodies have eventually failed to neutralize the Omicron variant [31].

While many clinical antibodies are used to neutralize other pathogens, treat autoimmune diseases, and combat cancer, there is an urgent call to prepare efficacy-improving strategies in advance or in early response phase before therapy resistance develops. Similar to our machine learning strategies, a few methods have been proposed: (1) to efficiently select broadly neutralizing antibodies based on the structure information of specific epitope [32], and (2) to identify antibody variants of high potency using the evolutionary information of antibody sequences alone which are encoded in generic protein language models [33]. Beyond these powerful methods, we emphasize the urgency of advanced preparation and early response for therapeutic antibody optimization without timely availability or abundance of experimental data, such as epitope characterization, structure determination [29], or activities of antibody variants [30]. Also, compared to the single substitutions of any given antibody are selected and combined [30], our approach combines with predictions of the neutralization activity landscapes for a large and diverse set of IgG antibodies. The success of activity landscape prediction was largely attributed to a novel antibody language model that, despite being 10-100 times smaller, outperformed competing protein language models thanks to protein pretraining - antibody fine-tuning, VH-VL cross-attention, and CDR masking. Future therapeutic development strategies shall further integrate computational and experimental approaches for synergistic accuracy and efficiency to design and re-design clinically effective antibodies and other drugs against quickly evolving targets, such as viral pathogens and oncogenic antigens.

## Methods and Materials

### Sex as a biological variable

Human blood specimens were collected from both male and female patients. Additionally, our study examined male and female mice, and similar findings are reported for both sexes.

### Human subject study and biosafety approvals for blood draws

Human blood specimens from convalescent COVID-19 patients and related research activities were implemented under NIH guidelines and the protocols approved by the Northwestern University Institutional Review Board (STU00205299) as well as the Institutional Biosafety Committee for COVID-19 research.

### B-cell sequencing (VDJ), bioinformatic analysis

B cells were isolated from blood of convalescent COVID-19 donors using the EasySep B kit (Stemcell Technologies, cat no 17954). SARS-nCoV-2 Spike RBD (Raybiotech, catalog no. 230- 30162) was biotinylated via a commercially available kit (Thermo Sceintific cat no. 21330) and the resulting biotin-labeled protein was purified through a Zeba quick spin column (Thermo Scientific cat no. 89882). The RBD-biotin was prebound with streptavidin-AlexaFluor-647 to make RBD-647 (Invitrogen, cat no. S21374). B cells were stained with CD19, CD27, CD38, anti-IgM and RBD-647 and a TotalSeq-C hashtag oligo (Biolegend). RBD-647+ B cells were sorted on a FACS Aria (BD Biosciences) in the Robert H. Lurie Cancer Center Flow Cytometry core facility. Due to the small number of RBD-647+ B cells isolated from each patient, we combined B cells with monocytes isolated from the same patients, labeled with a different hashtag oligo (24) .

B Cells and monocytes were partitioned using a 10X genomics Chromium Controller for GEM generation followed by single-cell library construction using 10X Chromium Next GEM Single Cell V(D)J Library reagent kit. Libraries were sequenced at the Northwestern University Sequencing Core Facility on an Illumina HiSeq 4000 using the sequencing parameters indicated by the manufacturer. We aligned the sequences aligned using CellRanger (10X genomics). Aligned antibody sequences from CellRanger were extracted for downstream analysis. VDJ sequences were ordered from IDT and cloned into the vector AbVec antibodies (Addgene)

### WT-neutralization prediction based on antibody structure prediction and docking

From each antibody sequence, we first predicted its 3D structure using the web application ABodyBuilder-ML (38). Then we determined the “active” residues for each antibody’s predicted structure using the web server proABC2 (39) and the “passive” residues for an RBD crystal structure (PDB ID: 6W41, Chain C) using the criteria of DSSP-calculated relative solvent accessibility above 0.4. For each pair of antibody heavy (H) and light (L) chains and RBD antigen structures, we performed initial protein docking using the webserver HADDOCK (25) (“Scenario 2” when a loose definition of the epitope is known) as well as the aforementioned information about active and passive residues. And we refined the resulting top-10 or less HADDOCK structural models (cluster representatives) and estimated their confidence weights using the computer program Bayesian Active Learning (BAL) (29). For each antibody, we calculated the confidence indicator of WT virus neutralization probability (0.0-1.0) using these protein-docking models to calculate antibody-blocking RBD residues from ACE2 binding. A total of 26 ACE2- binding residues on RBD were determined using the ACE2-RBD co-crystal structure (PDB ID: 6M17, 6M0J, and 6LZG), based on the proximity within 5Å of the ACE2; their residue indices are 417, 446, 447, 449, 453, 455, 456, 473, 475, 476, 477, 484, 486, 487, 489, 490, 493, 494, 495, 496, 498, 500, 501, 502, 503, and 505. Specifically, each given antibody is predicted to interact and interfere with a portion (0.0-1.0) of the 26 ACE2-binding residues on RBD(within 5Å of any RBD heavy atom) according to each structural model; and the WT-neutralizing confidence predictor is calculated by weight-averaging the portions across all structural models for each antibody.

### Antibody language model (AbLM)

The architecture, training, and application of AbLM is illustrated in **Fig. 2a**. We adopted a bidirectional self attention-based transformer encoder (12 layers and 12 heads per layer; please refer to the model ‘RP15_B1’ in (40) for more details). We pretrained the encoder using over 12 million non-redundant protein-domain sequences from Pfam-RP15-v32 (41) and fine-tuned two weight-tied sequence encoders, one for the heavy chain (VH) and the other for the light chain (VL), using 4,196 paired heavy and light-chain antibody sequences (variable region) from SabDab (42). During fine-tuning, a VH-VL cross-attention module was added after the weight-tied VH and VL encoders. While random masking following BERT was applied in pre-training, antibody-specific masking of one random CDR region per training example was adopted in fine-tuning. For the input of given paired heavy and light-chain sequences, the output of the resulting antibody language model is the “embedding” of the antibody sequence, that is, a 1536-dimensional vector consisting of 768 dimensions for either heavy or light chain. The embeddings lie in a 1536- dimensional space dubbed the “latent” space.

### Variant response prediction based on machine learning

To emulate the low-data regimes, we used variant response profiles of 14 clinical antibodies as of late 2022 (including 9 single antibodies under emergency use authorization and 5 single antibodies under clinical trials), as in “susceptibility summaries” of the Coronavirus Resistance Database (CoV-RDB) (43)) and **Supp. Fig. S6**. Specifically, we used their log fold improvements against Delta, Omicron/BA.1, and Omicron/BA.4/5, ranging between -3 (≥1000-fold reduced neutralizing activity) and +1 (≥10-fold increased neutralizing activity). In the 1536-dimensional latent space, we constructed covariance kernels using Euclidean distances among the embeddings of 1366 uncharacterized IgG candidates and 14 experimentally profiled clinical antibodies and accordingly three Kriging regressors for log fold improvements against Delta, Omicron/BA.1, and Omicron/BA.4/5, respectively. We made variant response predictions for all 1366 IgG candidates and visualized the landscape by interpolation (inverse distance weighting). We then validated such prediction based on 19 IgG candidates.

### Antibody re-design against the Delta variant

We chose antibody 106/107 as the “seed” for improved neutralization against a SARS-CoV-2 variant (Delta). The highest-weight structural model of the antibody-RBD (WT) complex as the input, a computational protein design program iCFN (36) is used to first predict the antibody-RBD (Delta variant) complex structure and then design amino-acid substitutions for the antibody to gain neutralization against the Delta variant. Specifically, both steps involve multistate design with single substrate per state (see more details in (36)) and find the optimal structures (and sequences, when applicable) to minimize the energy difference between a positive and a negative state.

When predicting the antibody-RBD (Delta variant) complex structure, we maximally disrupt binding by minimizing the folding-energy difference between the separate (positive-state) and bound (negative-state) antibody-RBD, as detailed in (44). All residues within 5Å of RBD residues L452 and T478 were treated flexible during design, while amino-acid substitutions L452K and T478R were enforced.

When designing the optimal single amino-acid substitutions for the antibody 106/107 and simultaneously predicting the structures, we maximally enhance RBD-binding by minimizing the folding-energy difference between the bound (positive-state) and the separate (negative-state) antibody-RBD while constraining the folding stability, as detailed in (36) (XRCC1 design). The antibody redesign positions are all residues within 5Å of RBD residues 452 and 478, including 1 on the heavy chain near residue 452 of RBD and 7 on the light chain near residue 478 of RBD.

### Cloning of antibody expression plasmids

The VDJ sequences of heavy and light chains of prioritized antibody pairs were obtained from the single cell VDJ sequencing. DNA fragments harboring the specific variable sequences were then synthesized by gBlock Gene Fragment technology by Integrated DNA Technologies (IDT). Synthesized double stranded DNA fragments were then cloned into AbVec2.0-IGHG1 (Addgene #80795), AbVc2.0-1.1-IGKC (Addgene #80796), and AbVec1.1-IGLC2-Xho1 (Addgene #99575) to generate the heavy chain, kappa light chain, and lambda light chains, respectively. The IgG expression vectors and protocols were generously shared by Drs. Jenna Guthmiller and Patrick Wilson at the University of Chicago. Proper insertions of cloned DNA fragments were confirmed by Sanger sequencing.

### Antibody production and purification

Plasmids expressing the heavy and light chain of each tested antibody were transfected into HEK293T cells (ATCC CRL-3216) by a calcium chloride transfection method. Briefly, 60-80% confluent HEK293T cells were transfected with about 19 micrograms of each heavy and light chain expression plasmids (up to 1044 µL nuclease free H2O) mixed with 188 µL of 2M CaCl2. Followed by dropwise addition of 1.25ml of 2X HBS buffer pH 7.12 (50 mM Hepes Acid, 280mM NaCl, 1.5 mM NA2HPO4) while introducing bubbles to the mix. Following incubation for 20 minutes at room temperature, 15.5 mL cDMEM was added to the mix and subsequently transferred to cells and incubated at 37°C 5% CO2 for 16 hours, after which cells were washed with 1X PBS and fresh cDMEM was added to cells and incubated for three more days. Supernatants were collected and spun at 2,000 x g for 10 minutes at 4°C to pellet cell debris. Antibodies were purified from culture supernatants by protein A agarose beads (Thermo Fisher PierceTM Protein A Agarose, 20333). Briefly, cell culture supernatant was added to pre-washed beads (500 µL) and incubated on a table top rocker for 2 hours at room temperature followed by overnight incubation at 4°C. Bead-bound antibodies were collected by centrifugation at 1800 xg for 10 minutes at 4°C (Break off) and washed in 1M NaCl followed by two 1X PBS washes. Antibodies were eluted by adding 3 ml 0.1M glycine-HCL (pH 7.12) and rocking at room temperature for 10 minutes. Following centrifugation at 1800 x g for 10 minutes at 4°C, supernatants containing the antibodies were neutralized by adding 200ul 1M Tris-HCl (pH 8.8). Antibody solutions were then concentrated using an Amicon protein concentrator (4 mL capacity, 30 kDa MWT cutoff) and buffer exchanged with 1X PBS and antibodies were stored at 4C.

### Quantification of antibody concentrations using ELISA

The Human IgG ELISA Kit (ab195215) was used to quantify the concentration of purified IgGs following the manufacturer’s instructions and optical density was measured using the SpectraMax iD5 plate reader.

### Cell-based neutralization assay

To create neutralized Spike Receptor Binding Domain (RBD), purified antibodies were incubated with the RBD-biotin-AF647 bait (3.3 nM) for 45 min on ice, then incubated with ACE2 expressing HEK-293 (ACE2^+^ HEK-293) cells (200,000 cells in 100 µL 2% extracellular vesicles (EV)-free FBS/PBS) for 45 min on ice. RBD bait that was incubated with PBS, or with non-fluorescent RBD bait (mock control) were used as controls. Cells were then washed twice with 2%EV-free/PBS (300xg for 5min). Dapi was added to stain dead cells and analysis was performed using BD FACSymphony A5-laser analyser. Viable singlets were gated for percentage of the RBD-AF647+ population. Data were analyzed using Flow Jo v10.6.2. and GraphPad Prism 9.5.0

### Live SARS-CoV-2 virus infection of A549 cells (BSL3)

The live virus neutralization experiments were conducted at the NIAID-supported BSL-3 facility at the University of Chicago Howard T. Ricketts Regional Biocontainment Laboratory.

One day prior to viral infection, A549 cells overexpressing human ACE2 (A549-hACE2) (obtained from tenOever and colleagues) (45) were seeded onto 96-well plates at a density of 10,000 cells per well. Antibody dilutions were made in Infection Media with 2% FBS. Antibody dilutions were mixed with 500 pfu of a SARS-CoV-2 strain, WT Washington (nCoV/Washington/1/2020), variant Delta (NR-55672 SARS-Related Coronavirus 2, Isolate hCoV-19/USA/MD-HP05647/2021 (Lineage B.1.617.2), variant Omicron (NR-56481 SARS-Related Coronavirus 2 Isolate hCoV- 19/USA/GA-EHC-2811C/2021 (Lineage B.1.1.529) or variant Omicron Isolate hCoV- 19/USA/COR-22-063113/2022 (Lineage BA.5) and incubated at 37°C in the dark for 1 hour. Culturing media was then replaced with infected antibody dilutions and allowed to incubate for 72-96 hours or until positive control wells show at least 50% CPE (cytopathic effect). At which point the infectious media was removed and cells were fixed with 100 µL of Formalin solution (10% Formalin Fisher 23305510). Cells were incubated at room temperature with Formalin for at least 15 minutes to ensure viral inactivation. Then formalin media was removed, and cells were stained in 0.25% Crystal Violet solution (0.25% w/v Crystal Violet Sigma C0775 in 20% EtOH) for 15-30 minutes, after which the Crystal violet is washed off under gently flowing tap water. Plates were then allowed to dry uncovered on the benchtop for at least 24 hours prior to analysis using the Infinite 200 Pro TECAN plate reader. Control wells such as no virus mock control in addition to a non-antibody treated control were included. A value of maximal death caused by the virus was evaluated from the virally infected but non-treated control wells while the maximal normal cell growth was determined from the mock-infected control wells. The average absorbance value of the non-treated control wells was subtracted from the remaining absorbance values to establish “0” values for non-treated wells. Next, all absorbance values were divided by the average absorbance value of the mock-infected control thereby setting the value of the mock-infected control to 100.

### Mouse experiments

Three experimental groups of 15 week-old hACE2 transgenic B6.Cg-Tg (K18-ACE2) 2Prlmn/J (K18-hACE2) mice were set up with nine mice per group (4 or 5 females and males each group to reach about 50%/50%). Mice were housed in specific pathogen-free facilities at the BSL-2 facility at University of Chicago Howard T. Ricketts Regional Biocontainment Laboratory. Three groups of IgG (1.2ug for Washington and Delta or 8.5ug for Omicron) were each mixed with SARS-CoV-2 viral strains; Washington at 10,000 pfu, Delta at 10,000 pfu, or Omicron at 20,000 pfu and incubated for 1 hour at 37°C. After incubation, the mice were anesthetized with isoflurane and 25 µL of the mixture were administered intranasally. Following viral infection challenges, animals were monitored for health twice daily and weighed once per day. Clinical scoring system included: Score 0 (pre-inoculation)- mice are bright, alert, active, normal fur coat and posture. Score 1 (post-inoculation, pi)- mice are bright, alert, active, normal fur coat and posture, no weight loss. Score 1.5 - mice present with slightly ruffled fur but are active OR weight loss might occur but does not reach 2.5%; recovery can be expected. Score 2 (pi)- ruffled fur OR less active OR < 5% weight loss; recovery might occur. Score 2.5 (pi)- ruffled fur OR not active but movies when touched OR hunched posture OR difficulty breathing OR weight loss 5-10%; recovery is unlikely but still might occur. Score 3 (pi)- ruffled fur OR inactive but moves when touched OR difficulty breathing OR weight loss at 11-20%; recovery is not expected. Score 4 (pi)- ruffled fur OR positioned on its side or back OOR dehydrated OR difficulty breathing OR weight loss > 20% OR labored breathing; recovery is not expected. Score 5 (pi) - death. Three days post infection, mice challenged with the Omicron variant were euthanized while the mice challenged with Washington and Delta variant were euthanized on day seven post infection. One lung was used to determine SARS CoV-2 viral genome levels and the other lung was fixed with formalin.

### Lung fixation and histopathological analysis

Lung tissue was submerged in 1 mL formalin for 48 hours. Formalin was removed and 1 mL formalin was added and incubated for an additional 12 days. Tissue was tested for viral inactivation and released. Fixed lungs were then embedded in paraffin and sectioned by routine procedures followed by H&E staining. Stained slides were scanned then analyzed using the NDP view2 software. Double-blinded evaluation of the percent of total lung surface area involvement was performed by a pathologist following a graded scheme adopted from a previous report (46).

### Lung viral genome level measurement using qRT-PCR

RNA was extracted from mouse lungs using the Nucleospin 96 RNA extraction kit as per written instructions (Macherey-Nagel 740709.4). Prior to being run through the binding columns, the lung tissue was collected in the R1A buffer provided with the kit and homogenized using a FastPrep fp120 homogenizer with 1.4 mm ceramic beads (Omni international SKU 19-645). RNA was eluted in RNAse free water and used for qRT-PCR. Which was performed using an Applied BioSystems Step One Plus Realtime PCR system, using the SupperScriptIII Platinum One-Step qRT-PCR Kit with ROX (Invitrogen 11745-500). Sample RNA was measured using a standard curve made by extracting viral RNA from lab viral stocks. The quantity of RNA in the viral stock was measured with a Nanodrop 2000. CDC recommended N2 primers and probe used in the qRT-PCR were purchased from IDT (10006824, 10006825, and 10006826).

## Acknowledgements

We are most grateful to Drs. Patrick Wilson and Jenna Guthmiller at the University of Chicago for sharing their antibody generation protocols and IgG constructs. We thank Dr. Dominique Missiakas at the University of Chicago for assistance with A/BSL3 research at the Howard T. Ricketts Laboratory (BSL3 management and practice of infectious disease research core, NIAID grant UC7AI180312). We are also thankful to the team of Northwestern COVID-19 Antibody and Cancer Collaborative Group and advisory members, especially Drs. Alfred L. George Jr., Leonidas C. Platanias, Rex L. Chishom, and William A. Muller for their scientific input and resourceful support for the project. The work was partially funded by the National Institute of General Medical Sciences (NIH/NIGMS) R35GM124952 (Y.S), the National Institute of Allergy and Infectious Diseases (NIH/NIAID) R01AI167272 and Chicago Biomedical Consortium Accelerator Award A-017 (H.L. and D.F.), Northwestern University Feinberg School of Medicine Emerging and Re-emerging Pathogens Program (EREPP) (H.L. and D.F.), and the R.H. Lurie Comprehensive Cancer Center Blood Biobank fund and Northwestern University NUCATs grant UL1TR001422 (M.I.). We gratefully acknowledge the support from the Northwestern University NUseq Core Facility and the RHLCCC Flow Cytometry Facility (Cancer Center Support Grant NCI CA060553). Flow Cytometry Cell Sorting was performed on a BD FACSAria SORP system and BD FACSymphony S6 SORP system, purchased through the support of NIH 1S10OD011996-01 and 1S10OD026814-01. Portions of this research were conducted with the advanced computing resources provided by Texas A&M High Performance Research Computing.

## Author contributions statement

H.F.A, W.T., A.D.H, J.W. co-led the bench experiments and computational analyses, prepared figures, and contributed to the manuscript writing. L.E., J.R.S, Y.Sun. participated in the studies and contributed to Figure preparation and writing. N.K.D., B.S., Y.J., R.I., Y. X., V.N., D.E., G.C.R., M.J.S., S.S. provided technical support or conducted bench experiments, and analyzed data. M.G.I. provided critical clinical resources and supervised the research project. H.L., D.F., and Y.S. designed experiments, analyzed data, supervised the work, and wrote the manuscript.

## Data availability

Data analyzed and generated are included in the supplementary tables.

## Code availability

Codes are available at https://github.com/Shen-Lab/COVID-Ab-LowDataRegime and https://zenodo.org/doi/10.5281/zenodo.10989934

