## Supplementary Figures for "Novel antibody language model accelerates IgG screening and design for broad-spectrum antiviral therapy"

### Supp. Figure S1

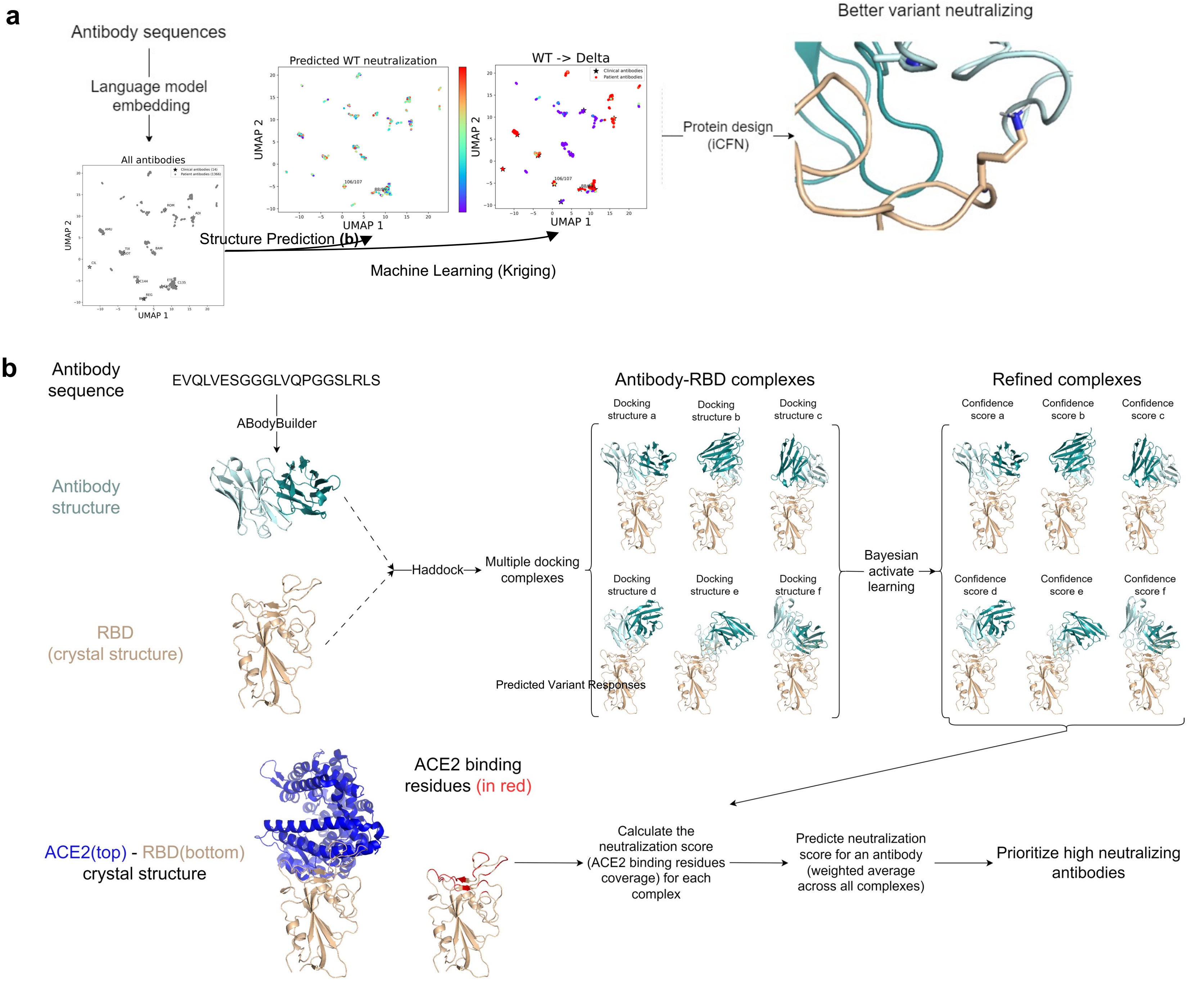


**Suppl. Figure S1. Schematics of the machine learning-assisted analysis of antibody sequences, structures, and activities.** (a) The overall pipeline consists of the following steps: embedding antibody sequences through our protein language model, predicting wild-type neutralization (by predicting antibody-antigen complex structures and the percentages of ACE2-binding residues of the RBD that are blocked by antibodies), and predicting variant responses (by Kriging of 14 clinical antibodies' known profiles in the language model-derived latent space). (b) The detailed steps of antibody-antigen structure prediction involve antibody 3D structure prediction through ABodyBuilder, antibody-RBD 3D structure docking through HADDOCK and BAL, and weighted neutralization score calculation.

#### Supp. Figure S2

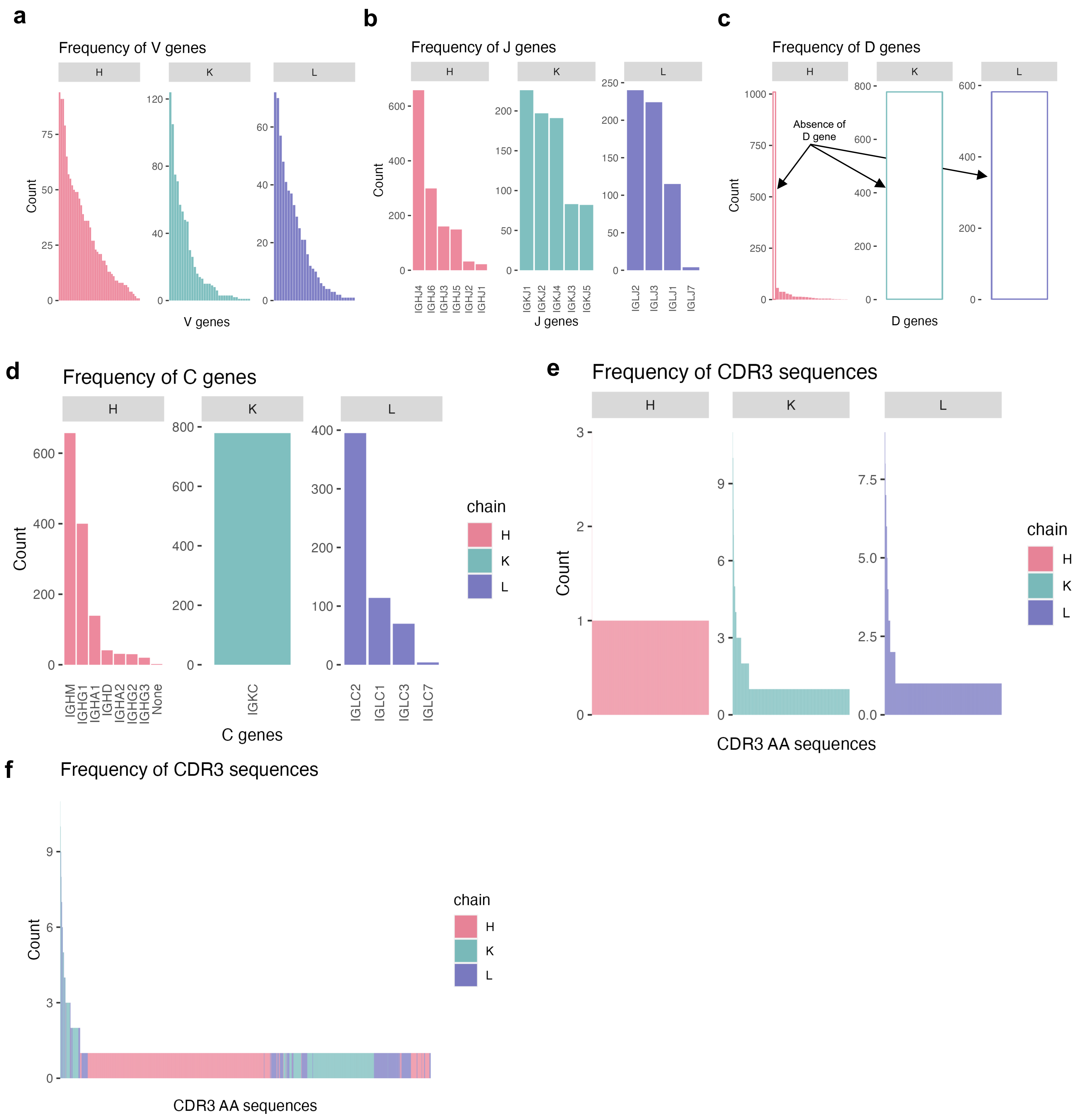


**Suppl. Figure S2. Frequencies of antibody C genes and CDR3 sequences.**
(a-c) Frequencies of V, J and D genes sequenced from 1366 RBD-bound B cells, within the antibody heavy (H) and Light chains (Kappa (K), and Lambda (L). (d) Frequency of C genes in H, K, and L chain sequences. (e, f) Frequency of CDR3 sequence in H, K, and L chains of VDJ sequences retrieved from the RBD-bound B cells of patients recovered from early wild-type SARS

### Supp. Figure S3

**a**

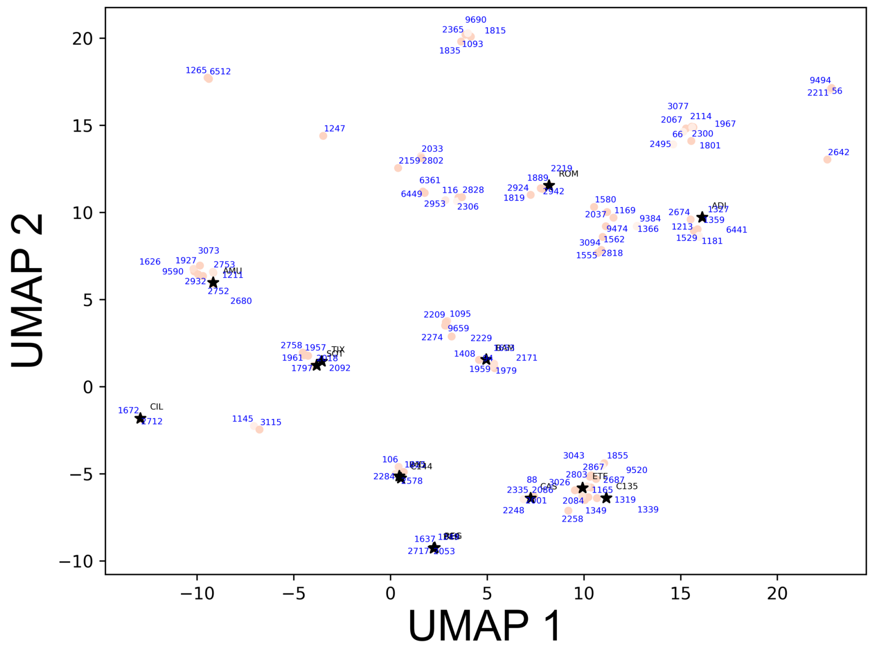


```
106_107H          1 EVQLVESGGGLVQPGGSLRLSCVASGITVSANYMSWVRQAPGKGLEWVSV     50
                    |||||||||||:||||||||||.|||.|||.||||||||||||||||||
C144H             1 EVQLVESGGGLIQPGGSLRLSCAASGFTVSNNYMSWVRQAPGKGLEWVSV     50

106_107H         51 IYSGGSTFYADSVKGRFTISRDNSKNTLYLQMNNLRADDTAVYSCAR---     97
                    |||||||:|||||||||||||||.||||||||||.|||:|||||.|||
C144H            51 IYSGGSTYYADSVKGRFTISRDKSKNTLYLQMNRLRAEDTAVYYCAREGE    100

106_107H         98 -----DF-----RGATAFDIWGQGTLVTVSS------------------    118
                         ||     |....||.|||||||||||
C144H           101 VEGYNDFWSGYSRDRYYFDYWGQGTLVTVSSASTKGPSVFPLAPSSKSTS    150


106_107L          1 QSALTQPASVSGSPGQSITISCTATSSDVDDYNYVSWYQQHPGKAPKLLI     50
                    ||||||||||||||||||||||||.|||||..|||||||||||||||||:|
C144L             1 QSALTQPASVSGSPGQSITISCTGTSSDVGGYNYVSWYQQHPGKAPKLMI     50

106_107L         51 YDVNNRPSGVSNRFSGSKSGNTASLTISGLQAEDEADYYCSSYTSSSTGV    100
                    |||:||||||||||||||||||||||||||||||||||||||||||||.|
C144L            51 YDVSNRPSGVSNRFSGSKSGNTASLTISGLQAEDEADYYCSSYTSSSTRV    100

106_107L        101 FGSGTKVTVL----------------------------------------    110
                    ||:|||||||
C144L           101 FGTGTKVTVLGQPKAAPSVTLFPPSSEELQANKATLVCLISDFYPGAVTV    150
```

**b**

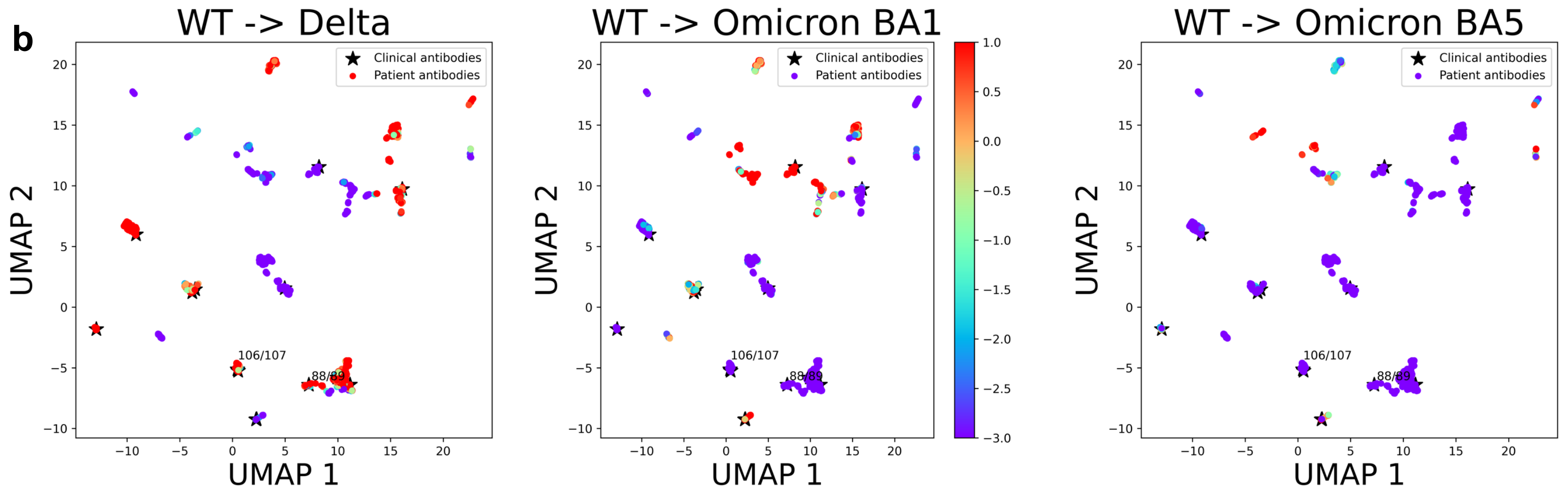


**Suppl. Figure S3. 2D visualization of the predicted patient activities in colors, where each antibody sequence is shown in the first two UMAP dimensions of the protein language model-derived 1536D space.** (a) A subset of 107 patient antibodies (in red dots and labeled with their heavy-chain indices) were predicted with high priority to neutralize the wild-type virus. Also shown is the sequence alignment between antibody 106/107 (H/L chains with CDR3 highlighted in yellow) and a clinical antibody CD144, its latent space neighbor.
(b) All 1366 patient antibodies were predicted with variant responses (log fold improvement compared to the wild type, ranging from -3 to +1). Also shown (in black stars) are 14 clinical antibodies whose variant profiles were used to predict the variant responses of patient antibodies.

#### Supp. Figure S4

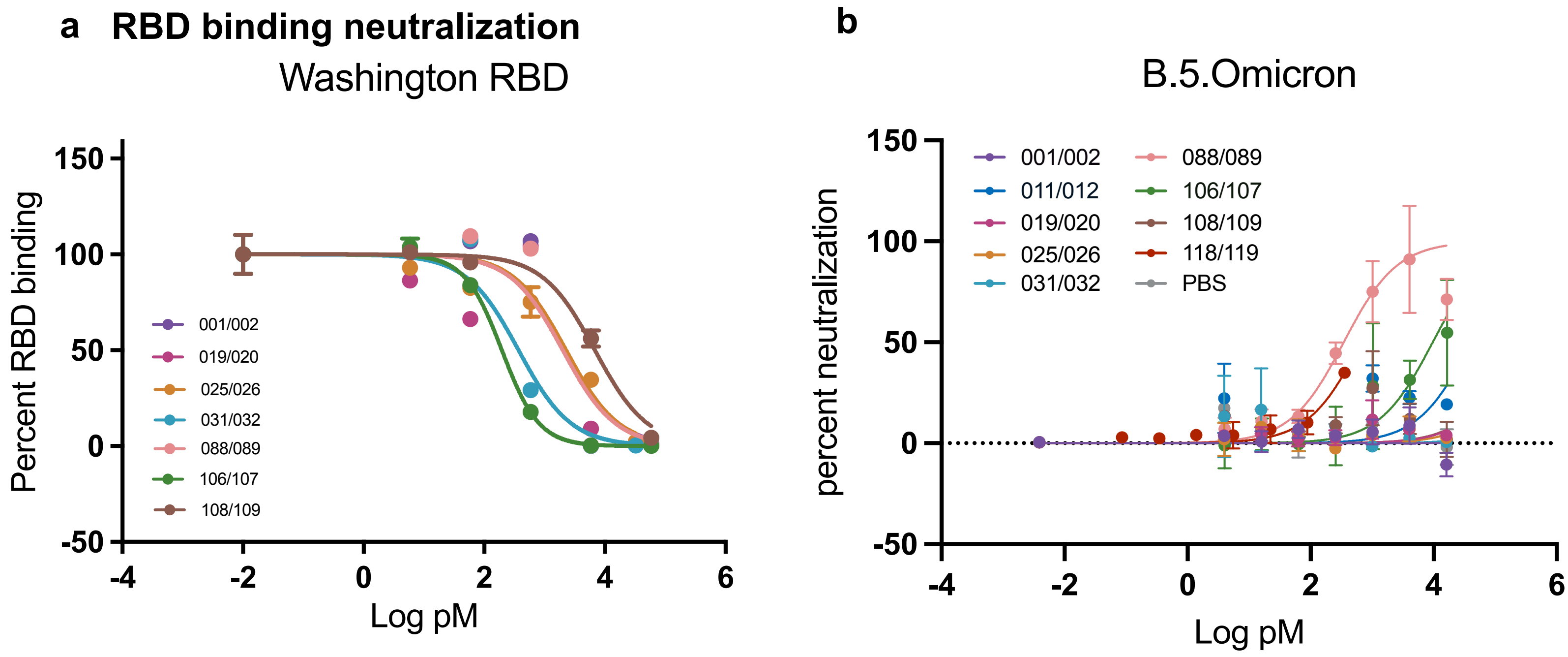


**Supp. Figure S4.** (a) Antibody neutralization of recombinant RBD in an ACE2+ HEK-293 cell-based neutralization assay. Increasing anti-RBD antibody concentrations were incubated with recombinant wildtype RBD followed by the addition of ACE+ overexpressing HEK-293T cells. The amount of non-neutralized RBD available to bind to ACE2 overexpressing cells was then followed using a flowcytometry approach. (b) Live SARS-CoV-2 virus neutralization by prioritized anti-RBD antibodies. Antibody dilutions were pre-incubated with 500pfu of SARS-CoV-2 Omicron B.5 variants. Following incubation, antibody-virus mix was added to A549 cells overexpressing human ACE2. At 96 hours post infection, cells were fixed, and cell viability was determined.

### Supp. Figure S5

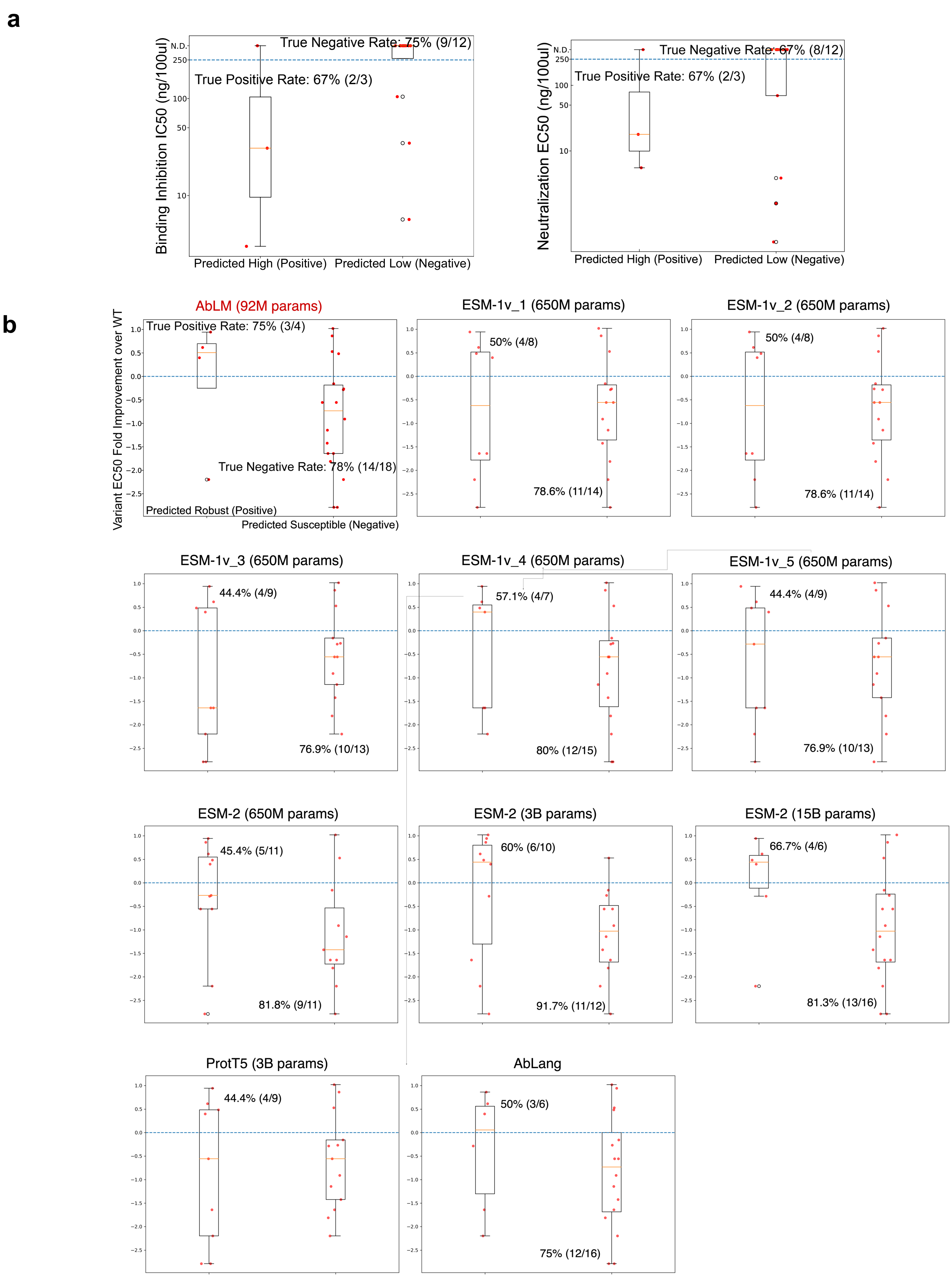

**Suppl. Figure S5. Assessing our predictions of IgG (a) wild-type viral neutralization and (b) variant susceptibility.** (a) Box plots of tested antibodies' wild-type neutralization (binding inhibition IC50 and viral neutralization EC50) split into three predicted to be of top/high-priority and twelve of bottom/low-priority. (b) Shown here are box plots of tested antibodies. (b) Performances of our novel Antibody Language Model (AbLM) versus state of the art protein language models (pLMs) in predicting antibody responses (robust or susceptible) to spike variants. variant response (log fold improvement in viral neutralization EC50 compared to the wild type) split into antibody-variant cases predicted to be robust or susceptible according to various pLMs, including our AbLM, five versions of ESM-1, ProtT5, three versions of ESM-2, and an antibody language model AbLang.  We note that AbLM had much higher precision for robust antibodies and better separation for robust versus susceptible antibodies, despite using 10-100 times less parameters.

### Supp. Fig S6

| Antibody | Delta | Omicron (BA1) | Omicron (BA5) | Omicron (BA2.75) |
|---|---|---|---|---|
| ADI | 1.5 | 108 | 935 | 257 |
| BAM | >1000 | >1000 | 686 | 424 |
| BEB | 1 | 1 | 1 | 10 |
| CAS | 0.7 | >1000 | >1000 | 384 |
| CIL | 2.1 | >1000 | 9.4 | 24 |
| ETE | 0.5 | 414 | 444 | 206 |
| IMD | 2.1 | >1000 | 633 | 384 |
| SOT | 1.3 | 3.8 | 16 | 9.1 |
| TIX | 1 | 306 | >1000 | 30 |
| REG | 28 | 1000 | 1000 | 42 |
| AMU | 0.6 | 136 | 116 | 57 |
| ROM | - | 0.8 | 64 | 5.3 |
| C135 | 0.4 | 1000 | 1000 | 1000 |
| C144 | 2.5 | 1000 | 1000 | 1000 |

**Suppl. Figure S6. Source data (14 clinical antibodies' known variant susceptibility from the Stanford CoVDB) that were used for predicting patient antibodies' variant responses.** Reported were fold changes in EC50.

#### Suppl. Figure S7

| Re-designed position [IMGT index (chain)] | Re-designed position [index in PDB files (chain)] | Proposed design (original amino acid -> proposed amino acid ) | △binding affinity for WT RBD (Kcal/mol; excluding VdW) | △binding affinity for delta RBD (Kcal/mol; excluding VdW) |
|---|---|---|---|---|
| 66 (H) | 57 (H) | PHE -> LYS | -0.59 | -3.24 |
| 66 (H) | 57 (H) | PHE -> ASN | -2.32 | -2.01 |
| 66 (H) | 57 (H) | PHE -> CYS | -1.65 | -2.71 |
| 38 (L) | 149 (L) | TYR -> SER | -9.9 | -11.51 |
| 38 (L) | 149 (L) | TYR -> ALA | -7.22 | -12.26 |
| 38 (L) | 149 (L) | TYR -> GLN | -8.35 | -11.82 |
| 38 (L) | 149 (L) | TYR -> GLY | -7.48 | -15.57 |
| 55 (L) | 166 (L) | TYR -> ASN | -5.94 | -5.02 |
| 55 (L) | 166 (L) | TYR -> ALA | -1.56 | -4.18 |
| 56 (L) | 167 (L) | L 167 ASP -> CYS | -1.49 | -4.04 |

**Supp. Figure S7. Computationally proposed top-designs (single amino-acid substitutions) of antibody 106/107 and their predicted binding-energy changes against the wild type or the Delta variant.** The highlighted designs Y38A (L) and Y38G (L), both hydrophobic substitutions, were predicted to stabilize binding to the Delta variant the most. They were both validated to improve neutralization of the Delta variant.

#### Suppl. Figure S8. Single variants

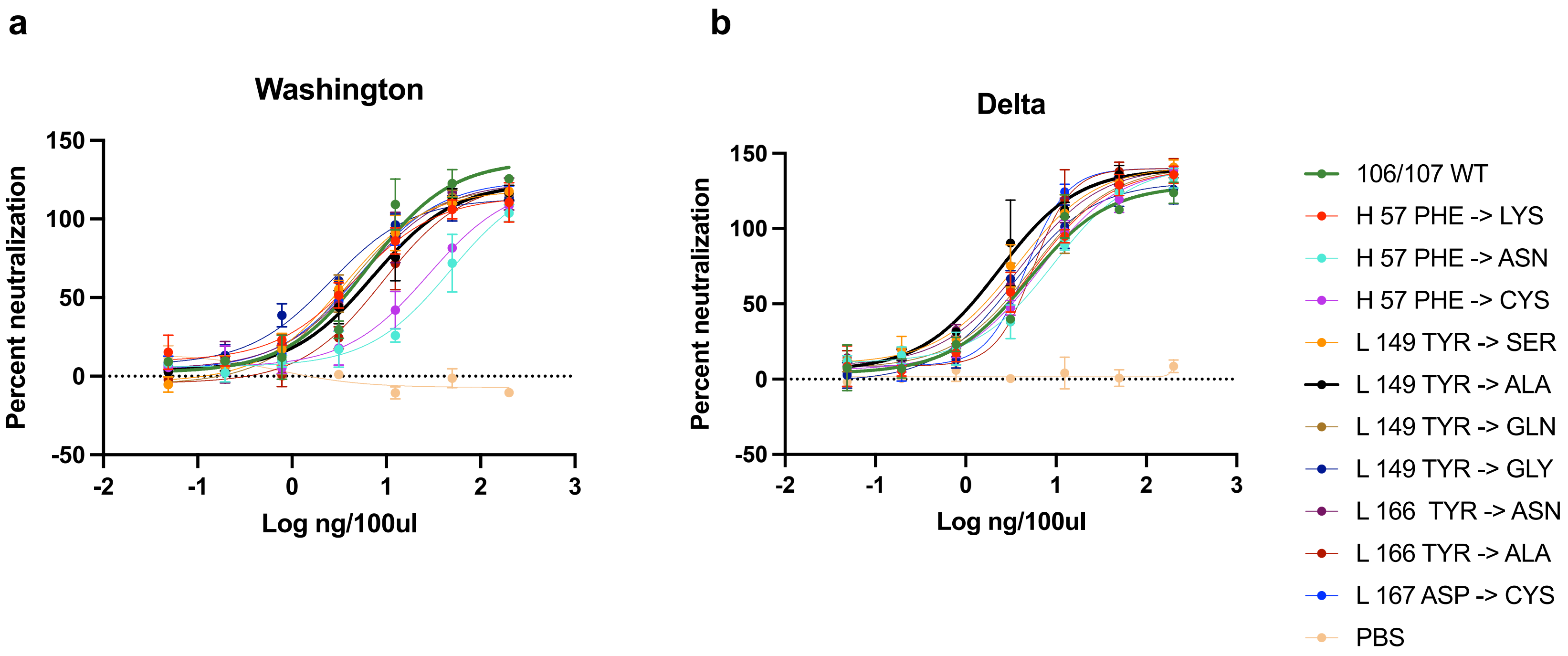


**Suppl. Figure S8.** Live SARS-CoV-2 virus neutralization by single variant mutated anti-RBD antibodies at increasing antibody concentrations. Percent neutralization of virus infecting A549 cells.

#### Supp. Figure S9

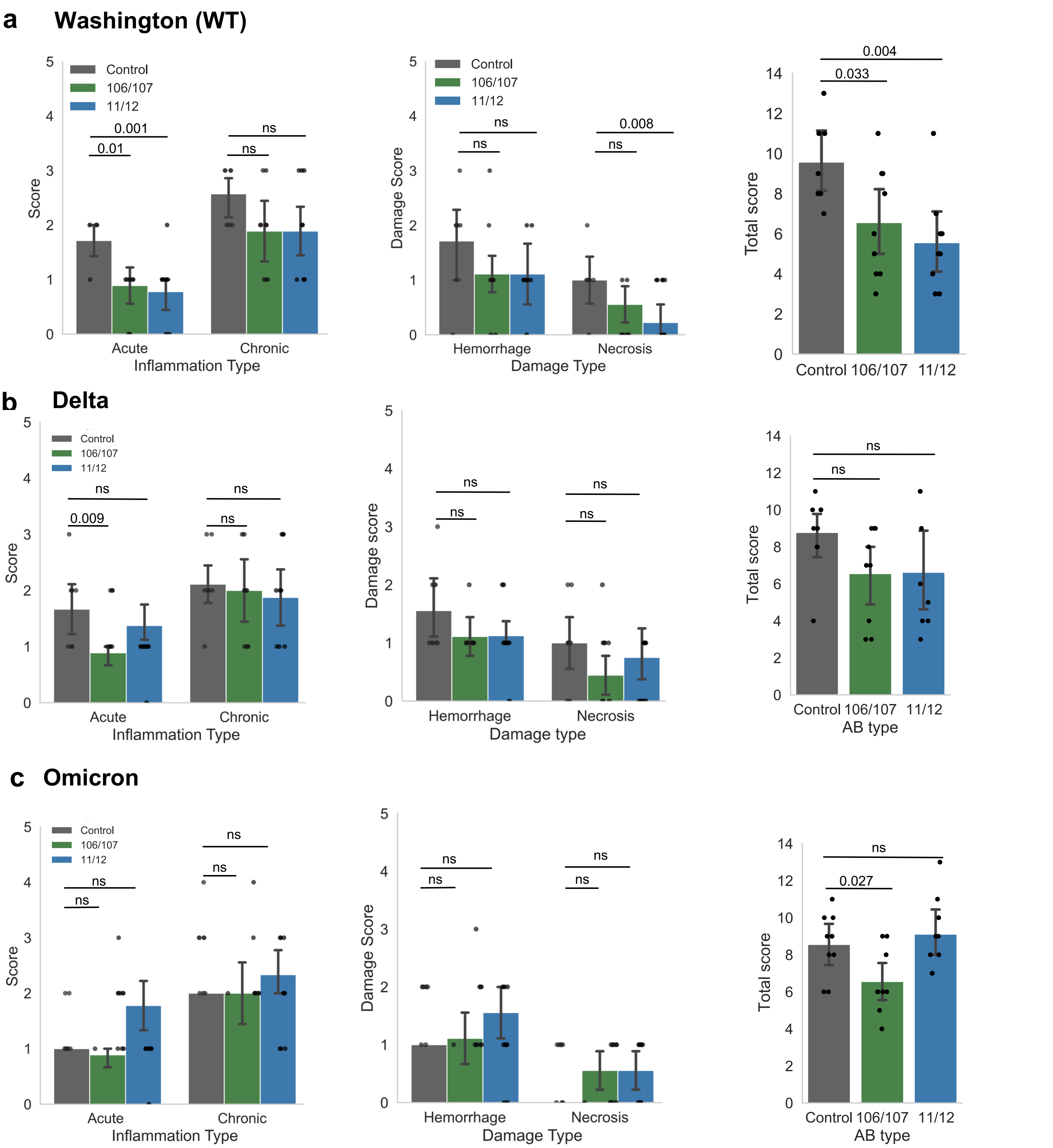


**Suppl. Figure S9. Lung histology inflammation and haemorrhage scores.** Alveolar hemorrhage and necrosis scores based on histopathological analysis of H&E-stained slides (n= 9). Statistical significance was tested using unpaired t-tests. ns means non-significant difference.

**Supp. Figure S10**

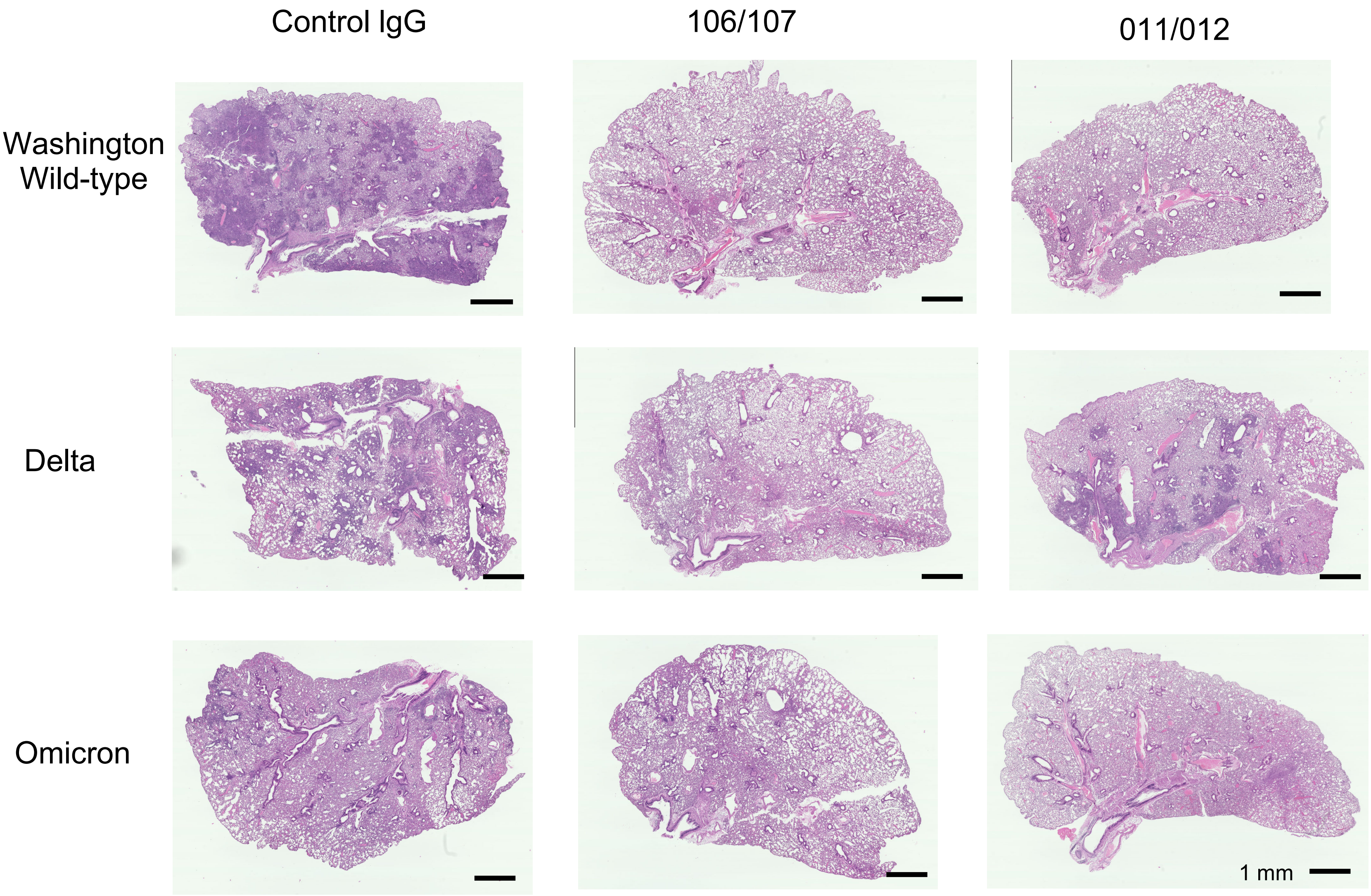


**Suppl. Figure S10. Lung histology images.** Representative H&E images of mice lungs at the experimental endpoint.
